# The furin cleavage site of SARS-CoV-2 spike protein is a key determinant for transmission due to enhanced replication in airway cells

**DOI:** 10.1101/2020.09.30.318311

**Authors:** Thomas P. Peacock, Daniel H. Goldhill, Jie Zhou, Laury Baillon, Rebecca Frise, Olivia C. Swann, Ruthiran Kugathasan, Rebecca Penn, Jonathan C. Brown, Raul Y. Sanchez-David, Luca Braga, Maia Kavanagh Williamson, Jack A. Hassard, Ecco Staller, Brian Hanley, Michael Osborn, Mauro Giacca, Andrew D. Davidson, David A. Matthews, Wendy S. Barclay

**Affiliations:** Department of Infectious Diseases, Imperial College London, UK, W2 1PG.; British Hearth Foundation Centre of Research Excellence, School of Cardiovascular Medicine & Sciences, King’s College London, UK, SE5 9RS; School of Cellular and Molecular Medicine, Faculty of Life Sciences, University of Bristol, UK, BS8 1TD; Department of Cellular Pathology, Northwest London Pathology, Imperial College London NHS Trust, UK, W6 8RF

**Author notes:** these authors contributed equally to this work (Thomas P. Peacock, Daniel H. Goldhill, Jie Zhou, Laury Baillon, Rebecca Frise).

## Abstract

SARS-CoV-2 enters cells via its spike glycoprotein which must be cleaved sequentially at the S1/S2, then the S2’ cleavage sites (CS) to mediate membrane fusion. SARS-CoV-2 has a unique polybasic insertion at the S1/S2 CS, which we demonstrate can be cleaved by furin. Using lentiviral pseudotypes and a cell-culture adapted SARS-CoV-2 virus with a S1/S2 deletion, we show that the polybasic insertion is selected for in lung cells and primary human airway epithelial cultures but selected against in Vero E6, a cell line used for passaging SARS-CoV-2. We find this selective advantage depends on expression of the cell surface protease, TMPRSS2, that allows virus entry independent of endosomes thus avoiding antiviral IFITM proteins. SARS-CoV-2 virus lacking the S1/S2 furin CS was shed to lower titres from infected ferrets and was not transmitted to cohoused sentinel animals. Thus, the polybasic CS is a key determinant for efficient SARS-CoV-2 transmission.

## Introduction

In 2019, a respiratory epidemic of unknown aetiology emerged in Hubei Province, China. The cause of the outbreak was quickly identified as a novel betacoronavirus, closely related to severe acute respiratory syndrome coronavirus (SARS-CoV) and named SARS-CoV-2 (P. Zhou et al., 2020; N. Zhu et al., 2020). SARS-CoV-2 is highly transmissible between humans and by the middle of March, the WHO declared the outbreak a pandemic (Holshue et al., 2020). It is vital to understand which molecular features of SARS-CoV-2 led to the virus causing a pandemic in order to control both the current pandemic and to prevent future coronavirus pandemics.

Coronaviruses enter host cells via their spike glycoprotein. Like many other enveloped virus glycoproteins, the spike protein is synthesised as a precursor that must be cleaved in order to perform its membrane fusogenic activity. Depending on the sequence of spike at the S1/S2 junction, the cleavage occurs during trafficking of spike in the producer cell by host furin-like enzymes, or by serine-proteases such as the transmembrane protease, serine 2 (TMPRSS2) at the cell surface after attachment, or by cathepsin proteases in the late endosome/lysosome (Matsuyama et al., 2010; Simmons et al., 2005). Upon cleavage of the S1/S2 junction and engagement of the host cell receptor with the spike receptor binding domain (RBD), a second cleavage site (CS) becomes exposed within the S2 domain, termed the S2’ site (Belouzard, Chu, & Whittaker, 2009; Millet & Whittaker, 2014). Upon cleavage of the S2’ site, generally by serine proteases or cathepsins, the S2 fusion peptide is liberated and initiates viral-host membrane fusion (Belouzard et al., 2009).

Like the closely related SARS-CoV, the cognate receptor of the SARS-CoV-2 spike is angiotensin-converting enzyme 2 (ACE2)(W. Li et al., 2003; P. Zhou et al., 2020). While the SARS-CoV S1/S2 junction is well characterised as being cleaved by serine proteases or cathepsins, the SARS-CoV-2 spike, similarly to the more distantly related Middle Eastern respiratory syndrome-related coronavirus (MERS-CoV), contains a polybasic CS, described as being a suboptimal furin CS (Coutard et al., 2020; Millet & Whittaker, 2014; Shang et al., 2020). This polybasic CS is absent from the closest relatives of SARS-CoV-2, such as SARS-CoV, although similar polybasic CS are found in more distantly related coronaviruses (Andersen, Rambaut, Lipkin, Holmes, & Garry, 2020; Boni et al., 2020; Le Coupanec et al., 2015). It has previously been demonstrated for the MERS-CoV spike, and for SARS-CoV-2, that the furin CS at the S1/S2 junction may promote entry into lung cells (Hoffmann, Kleine-Weber, & Pohlmann, 2020; Park et al., 2016). Interestingly, SARS-CoV-2 has been shown in multiple independent studies to rapidly lose this polybasic CS upon passage in Vero E6 cells, which has been a popular cell line for isolating and propagating the virus (Davidson et al., 2020; Klimstra et al., 2020; Z. Liu et al., 2020; Ogando et al., 2020; Sasaki et al., 2020; Wong et al., 2020; Y. Zhu et al., 2020). In addition, there are reports of CS mutants isolated directly from clinical swabs (Z. Liu et al., 2020; Wong et al., 2020). Several different mutants in this region are described including total deletions of the CS, loss of arginine mutations within the CS making it less polybasic, or deletions of flanking regions leaving the polybasic tract intact but potentially affecting accessibility to protease.

In this study, we use a combination of lentiviral pseudotypes with spike CS mutations and Vero passaged SARS-CoV-2 virus variants to investigate the molecular mechanism by which the polybasic CS of SARS-CoV-2 mediates efficient entry into lung cells. We describe the biological consequences of these mutations and test the effect of these mutations on viral transmission in ferrets.

## Results

### The polybasic S1/S2 cleavage site of SARS-CoV-2 spike protein is cleaved by furin

To investigate the importance of the spike polybasic CS of SARS-CoV-2 (PRRAR), a number of spike mutants were generated which were predicted to modulate the efficiency of furin cleavage (Figure 1A) including: substituting two of the upstream arginines to produce a monobasic CS similar to SARS-CoV spike (monoCS), replacing the tribasic CS with the furin CS of a highly pathogenic H5N1 avian influenza haemagglutinin containing a string of seven basic amino acids (H5CS), and two naturally occurring deletions seen following passage in Vero E6 cells and/or in clinical isolates (Davidson et al., 2020; Lau et al., 2020). The first of these removes eight amino acids including all 3 arginines of the PRRAR site (ΔCS) – while the other removes five flanking amino acids but retains the tribasic site (Δflank). The mutations were engineered into a cDNA encoding the spike to enable cell surface expression and the generation of lentiviral pseudotypes (PV) that carry each spike variant. In addition, to study the importance of the PRRAR motif in the context of live virus we took advantage of a naturally occurring Vero cell adapted mutant SARS-CoV-2 isolate, ΔCS (Davidson et al., 2020). This variant and the wild type virus from which it was derived were cloned by limiting dilution to enable studies using individual genotypes.

**Figure 1.**
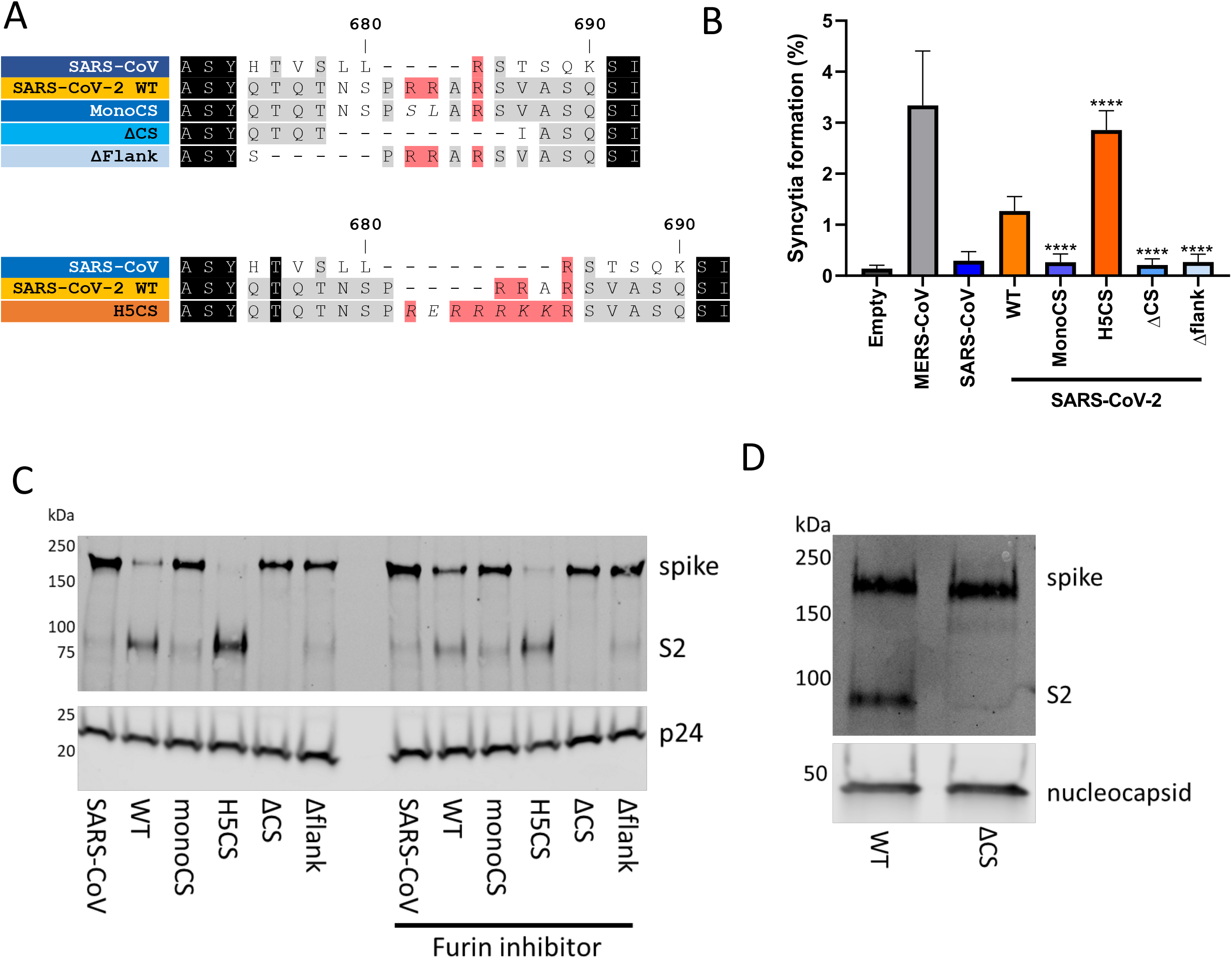
The SARS-CoV-2 spike contains a suboptimal polybasic furin cleavage site at the S1/S2 site. (A) Amino acid sequence alignment of coronavirus furin cleavage site mutants used in this study. Mutants with potential S1/S2 furin cleavage sites shown in shades of orange while mutants without furin cleavage sites shown in shades of blue. (B) Syncytia formation due to overexpression of different coronavirus spike proteins in Vero E6 cells. Percentage indicates proportion of nuclei in each field which have formed clear syncytia. Statistical significance determined by one-way ANOVA with multiple comparisons against SARS-CoV-2 WT. **** indicates P value < 0.0001. (C) Western blot analysis of concentrated lentiviral pseudotypes with different coronavirus spike proteins. Levels of lentiviral p24 antigen shown as loading control. Lentiviral pseudotypes labelled ‘furin inhibitor’ were generated in the presence of 5 μM Decanoyl-RVKR-CMK, added 3 hours post-transfection. (D) Western blot analysis of concentrated WT and ΔCS SARS-CoV-2 viruses. Levels of nucleocapsid (N) protein shown as loading control.

In several previous studies, the ability of coronavirus spike proteins to be cleaved by furin has been correlated with the ability to generate syncytia at neutral pHs when overexpressed (Belouzard et al., 2009; Hoffmann, Kleine-Weber, & Pohlmann, 2020; Xia et al., 2020). The library of SARS-CoV-2 mutant spike proteins were transiently expressed in Vero E6 cells, which do no express TMPRSS2 (Bertram et al., 2010; Shirato, Kawase, & Matsuyama, 2013), and syncytia formation was compared to SARS-CoV and MERS-CoV spikes (which have previously been shown to poorly and efficiently result in syncytia formation, respectively). As described before, SARS-CoV spike expression resulted in poor syncytia formation while MERS-CoV spike produced much higher levels of syncytia (Figure 1B). SARS-CoV-2 WT spike gave an intermediate level of syncytia formation that was ablated for the mutants which were not cleaved by furin. The H5CS spike bearing the optimised furin CS produced a higher level of syncytia formation than SARS-CoV-2 WT, similar to MERS-CoV spike.

To investigate the differences in spike cleavage efficiency in producer cells between the mutants, we produced PV with each mutant spike protein (or SARS-CoV) in human embryonic kidney 293T (293T) cells. PV were concentrated and probed by western blot (Figure 1C, left panel). Equal amounts of PV particles were loaded as indicated by p24 content. PV formed with SARS-CoV-2 WT spike had two bands reactive with anti-spike S2 antibody, corresponding to cleaved and uncleaved spike, with the stronger band corresponding to the cleaved S2 product. PV containing the H5CS spike showed very little uncleaved spike while PV with SARS-CoV WT spike and SARS-CoV-2 monobasic and deletion mutants showed near compete losses of cleaved spike. When PV were produced in parallel in the presence of a furin inhibitor, full-length spike was somewhat restored for the WT and H5CS spike (Figure 1C, right panel). We also took WT and ΔCS SARS-CoV-2 virus, concentrated virions by centrifugation and probed by western blot for spike cleavage (Figure 1D). Like the PV, WT SARS-CoV-2 harboured both uncleaved and cleaved S2 whereas the virions of the ΔCS mutant virus only had uncleaved spike. Overall, these data confirm that the polybasic CS of SARS-CoV-2 is a *bona fide* furin CS.

### The polybasic cleavage site of SARS-CoV-2 spike protein promotes entry into epithelial cell lines and cultures but adversely affects entry into Vero and 293Ts cells

To investigate whether the S1/S2 furin CS of SARS-CoV-2 plays a role in virus entry, we initially performed competition assays by taking a mixed population of virus containing 70% ΔCS mutant and 30% WT SARS-CoV-2 (as determined by deep sequencing of the S1/S2 CS; Figure 2A) and inoculating the virus mix onto Vero E6 cells, human intestinal Caco-2 cells or air-liquid interface human airway epithelial cell cultures (HAEs) at a low multiplicity of infection (MOI) to enable multicycle replication. We deep sequenced the progeny virus at 72 hours post-inoculation from the Vero E6 or Caco-2 (human intestinal) cells and found that whereas the ΔCS mutant outcompeted the WT in Vero E6 cells, WT became predominant in the Caco-2 cells. In primary HAE cultures, the WT virus also outcompeted the ΔCS virus until the variant was almost undetectable after 72 hours. We also infected Calu-3 (human lung) cells with either the clonal WT or ΔCS viruses at an MOI of 0.1 (Figure 2C). WT virus replicated robustly and reached peak titres greater than 10^5^ pfu after 48 hours. Conversely, ΔCS virus, appeared unable to productively infect Calu-3 cells and no infectious titre was detected in supernatant at any time point.

**Figure 2.**
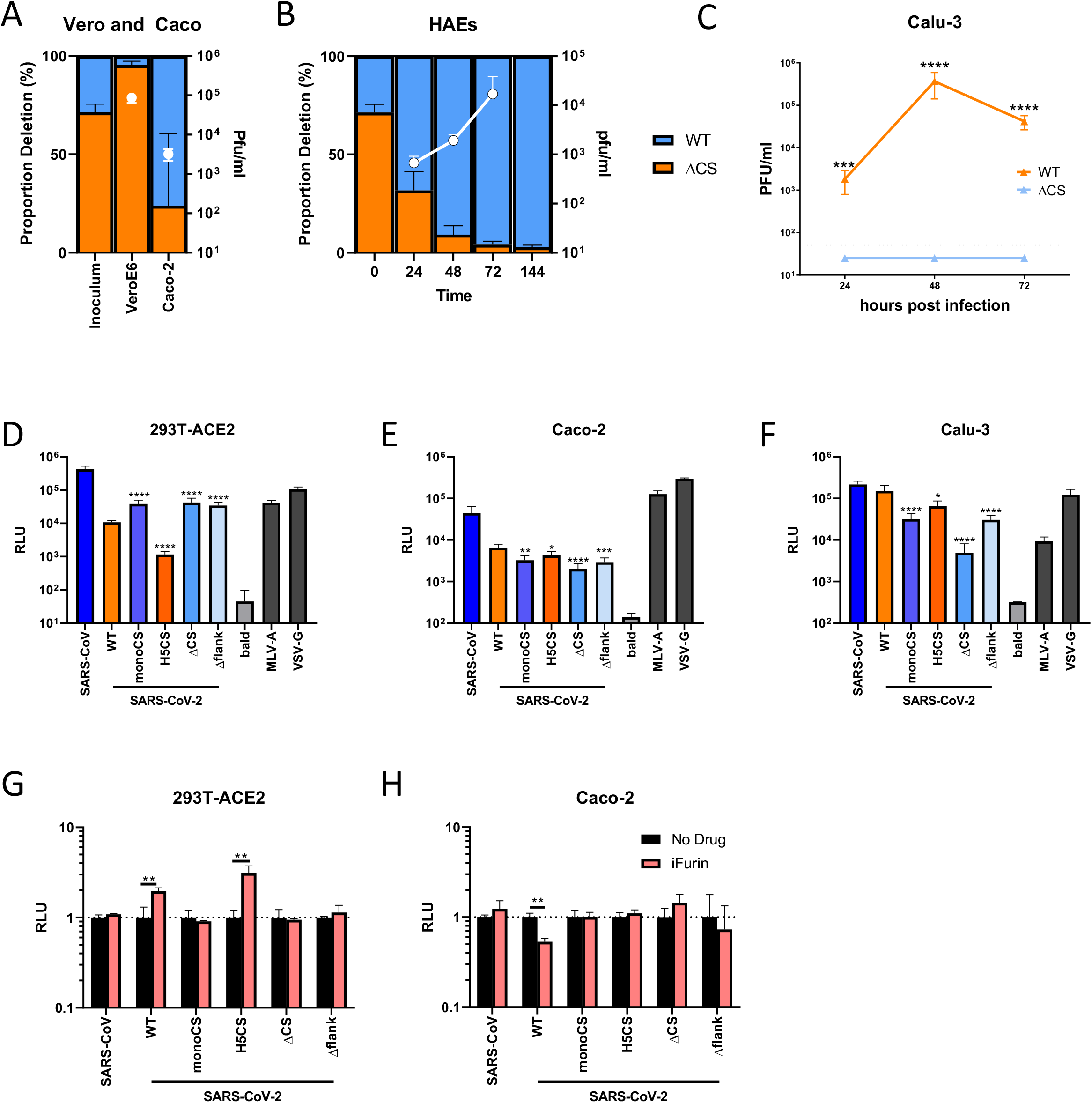
The furin cleavage site of SARS-CoV-2 mediates entry into mucosal epithelial and primary human airway cells. (A) SARS-CoV-2 competition assay growth curve between WT and ΔCS virus in Vero E6 and Caco-2 cells. Cells infected at an MOI of 0.1. Starting inoculum ratio shown on the left-hand bar while proportions of virus as determined by deep sequencing at 72 hours post-inoculation shown on the right. Virus titres determined by plaque assay at 72 hours post-inoculation shown in superimposed white data points. All results indicate triplicate repeats. (B) SARS-CoV-2 competition assay growth curve between WT and ΔCS virus in human airway epithelial cells (HAEs). Cells infected at an MOI of 0.1. Starting inoculum ratio shown at time 0, proportions of virus determined by deep sequencing. All time points taken from triplicate repeats. Virus replication determined by plaque assay and shown as imposed white data points. (C) Head to head replication kinetics of clonal WT and ΔCS viruses in Calu-3 human lung cells. Cells infected at an MOI of 0.1. All time points taken from triplicate repeats. Virus replication determined by plaque assay. Statistics determined by Student’s t-test on log transformed data. ***, 0.001 ≥ P > 0.0001; ****, P ≤ 0.0001. (D,E,F) Entry of lentiviral pseudotypes (PV) containing different viral glycoproteins into 293T-ACE2 (D), Caco-2 (E) and Calu-3 (F) cells. Cells transduced with different PV and lysed 48 hours later and analysed by firefly luciferase luminescence. All assays performed in triplicate. Statistics determined by one-way ANOVA on Log-transformed data (after determining log normality by the Shapiro-Wilk test and QQ plot.) *, 0.05 ≥ P > 0.01; **, 0.01 ≥ P > 0.001; ***, 0.001 ≥ P > 0.0001; ****, P ≤ 0.0001. (G,H) Relative entry of PV grown in the absence or presence of furin inhibitor (5 μM Decanoyl-RVKR-CMK) into 293T-ACE2 (G) or Caco-2 (H) cells. Untreated PV normalised to an RLU of 1. Statistics determined by multiple t-tests. All assays performed in triplicate. **, 0.01 ≥ P > 0.001.

Next, we probed the ability of PV with different mutant spike proteins to enter several different human cell lines: 293T cells expressing human ACE2, Caco-2 cells or Calu-3 cells (Figure 2D-F). PV bearing the envelope of amphotropic murine leukaemia virus (MLV-A) or Indiana vesicular stomatitis virus glycoprotein (VSV-G) were used as positive controls for cell entry while PV produced without any viral glycoproteins (bald) were used as negative controls throughout. As seen in the Vero E6 cells (Figure 2A), a clear negative correlation was seen between efficiency of furin cleavage of the spike and entry in the 293T-ACE2 cells (Figure 2D). PV with WT SARS-CoV-2 spike entered 293T-ACE2s more poorly than SARS-CoV, while mutants without furin cleavage (monoCS, ΔCS, Δflank) entered cells significantly more efficiently (over 3-fold compared to WT). Introduction of the optimal furin CS (H5CS) resulted in significantly poorer entry than WT (^~^10-fold lower; P < 0.001). When Caco-2 and Calu-3 cells were tested for PV entry, however, the opposite trend was observed reflecting the efficiency of virus replication in Caco-2, Calu-3 and primary HAE cells (Figure 2E,F). WT and H5CS spike PV entered cells efficiently while the mutants unable to be cleaved by furin, including ΔCS, entered cells significantly less efficiently (>2-fold lower in Caco-2 cells and ^~^5-fold lower in Calu-3 cells).

We next tested the PV which had been generated in the presence of a furin inhibitor for their ability to enter 293T-ACE2 and Caco-2 cells (Figure 2G,H). In 293T-ACE2, entry of PV bearing WT SARS-CoV-2 or H5CS spike was boosted 2- to 3-fold if furin was inhibited when the PV was produced (Figure 2G). Conversely, WT SARS-CoV-2 PVs entered Caco-2 cells more poorly (2-fold lower) after inhibition of furin cleavage, while no effect was seen on the H5CS mutant, potentially due to the majority of spike still remaining cleaved even after furin inhibition (Figure 1H). No significant differences in entry were observed for the other mutants or SARS-CoV.

Overall, these results suggest that during replication of SARS-CoV-2 in Vero and 293T-ACE2 cells, there is a fitness cost in having a cleaved spike prior to entry, while in primary airway cells and lung and intestinal cell lines, possessing a processed SARS-CoV-2 furin CS provides an advantage by facilitating entry.

### Entry of SARS-CoV-2 into 293T cells is dependent on cathepsins while entry into Caco-2, Calu-3 and primary HAE cells is dependent on TMPRSS2

As well as processing at the S1/S2 CS, coronavirus spike protein requires cleavage at the S2’ site to enable the virus membrane to fuse with the membrane of the host cell. To investigate whether the different cell entry phenotypes seen in 293T-ACE2/Vero vs Caco-2/Calu-3/HAE cells was due to differences in the protease use in different cell types, we performed PV entry assays in the presence of different protease inhibitors: camostat which inhibits serine proteases such as TMPRSS2, and E-64d, which inhibits cathepsins. Both drugs have previously been shown to be inhibitory to SARS-CoV and SARS-CoV-2 entry (Hoffmann, Kleine-Weber, Schroeder, et al., 2020; X. Ou et al., 2020).

In 293T-ACE2 cells, camostat pre-treatment did not inhibit PV entry whereas E-64d strongly inhibited entry of SARS-CoV spike PVs, as well as SARS-CoV-2 WT PVs and all CS mutants (Figure 3A). In Caco-2 cells, a different pattern was seen: camostat had a significant impact on PVs bearing spike proteins with furin CSs whereas E-64d had a significant impact on PV which were not cleaved by furin (Figure 3B). In Calu-3 cells, camostat significantly inhibited entry of all coronavirus PVs while E-64d also had a modest, but significant (P < 0.05), effect on the ΔCS mutant (Figure 3C). Control PV expressing MLV-A or VSV-G, which are not reliant on cathepsins or serine proteases for entry (Hoffmann, Kleine-Weber, Schroeder, et al., 2020), were not significantly affected by either drug in any cell line.

**Figure 3.**
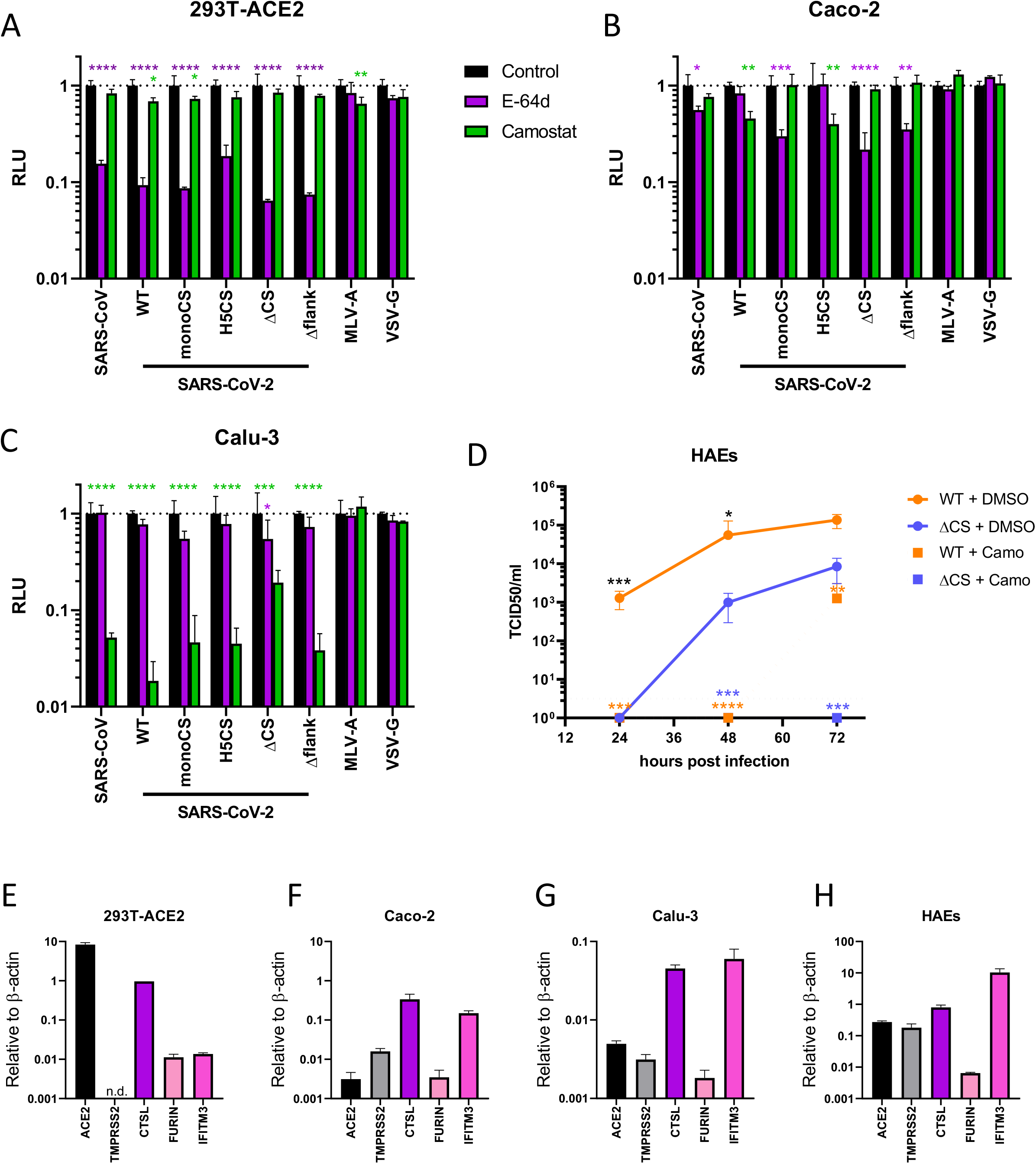
The furin cleavage site of SARS-CoV-2 spike allows more efficient serine-protease dependent entry into airway cells. (A,B,C) Inhibition of entry of lentiviral pseudotypes into (A) 293T-ACE2, (B) Caco-2 or (C) Calu-3 cells by the serine protease inhibitor, camostat (green bars) or the cathepsin inhibitor, E64-d (Purple bars). All data normalised to no drug control (black bars). Statistics determined by two-way ANOVA with multiple comparisons against the no drug control. *, 0.05 ≥ P > 0.01; **, 0.01 ≥ P > 0.001; ***, 0.001 ≥ P > 0.0001; ****, P ≤ 0.0001. (D) Replication kinetics of SARS-CoV-2 WT and ΔCS viruses in HAE cells. Cells were pretreated with control media or media containing camostat for 1 hour then infected at an MOI of 0.1. Statistics were determined by one-way ANOVA with multiple comparisons on log transformed data. Black asterisks indicate statistical significance between no drug controls of WT and ΔCS while coloured asterisks indicate significance between no drug control or camostat. *, 0.05 ≥ P > 0.01; **, 0.01 ≥ P > 0.001; ***, 0.001 ≥ P > 0.0001; ****, P ≤ 0.0001. (E-H) Gene expression of select SARS-CoV-2 entry factors in (E) 293T-ACE2, (F) Caco-2, (G) Calu-3 or (H) HAEs. Gene expression determined by qRT-PCR and normalised to *β-actin*

To confirm that dependence of serine proteases for entry, as observed in Caco-2 and Calu-3 cells, was seen with whole SARS-CoV-2 virus in primary airway cells, we examined multicycle replication of clonal WT and ΔCS viruses on HAE cells in the presence or absence of camostat (Figure 3D). Our previous results showed that the ΔCS virus was selected against on HAE cells and, indeed, the ΔCS virus grew to significantly lower titres than WT. Addition of 50 μM camostat almost completely abrogated virus replication of both ΔCS and WT virus (Figure 3D), showing that in HAEs, efficient cleavage by serine proteases facilitates efficient entry.

To investigate whether differences in endogenous levels of different proteases could explain the phenotypic differences seen in the different human cell lines, as well as HAEs, we quantified the expression of several different proteases using qPCR (Figure 3E-H). All three human cell lines and the primary HAE cultures expressed ACE2 and cathepsin L to varying degrees. However, 293T-ACE2 cells lacked any detectable TMPRSS2 expression, likely explaining why camostat had little effect on PV entry into these cells. Although not tested here, several previous studies have shown Vero E6 cells also express no endogenous TMPRSS2 (Bertram et al., 2010; Shirato et al., 2013).

### Expression of TMPRSS2 promotes entry of SARS-CoV-2 expressing a full polybasic cleavage site

To investigate whether expression of TMPRSS2 was responsible for the advantage seen by viruses with furin cleavable spike proteins in Caco-2 and Calu-3 cells, we transiently co-expressed ACE2 with or without TMPRSS2 in 293T cells and compared the entry of furin cleaved and non-furin cleaved PV (Figure 4A). We found that entry of all coronavirus PVs was promoted by the expression of TMPRSS2, even though TMPRSS2 expression led to lower levels of cell associated ACE2 due to ACE2 being a substrate of TMPRSS2 (Supplementary Figure S1A) (Shulla et al., 2011). The TMPRSS2 enhancement of PV entry was particularly potent for the PVs harbouring furin CS containing spike (>15-fold), compared to the non furin-cleaved mutants (<10-fold) indicating that expression of TMPRSS2 favours the entry into cells of PVs with furin CSs.

**Figure 4.**
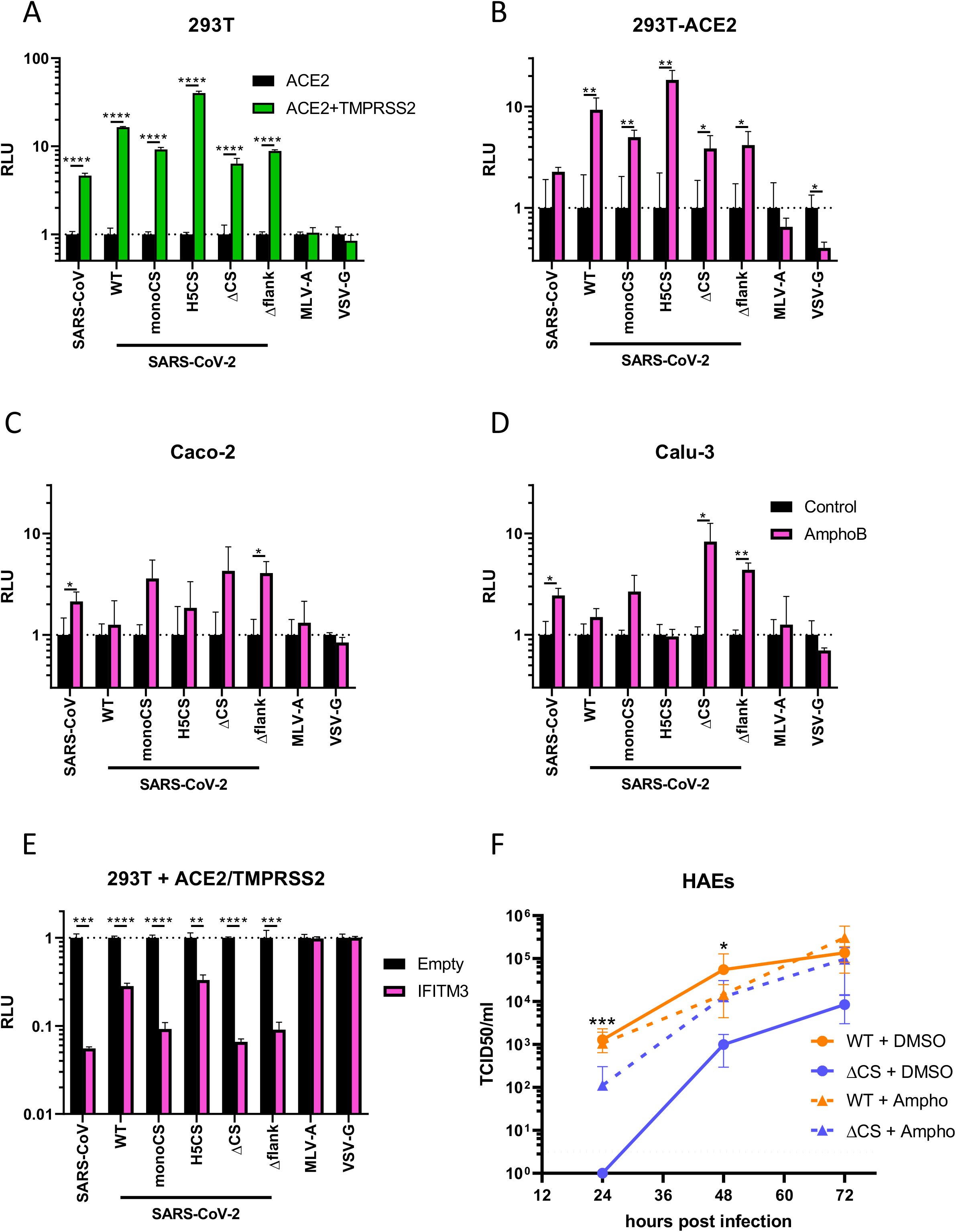
The efficient furin cleavage site-dependent entry of SARS-CoV-2 is due to TMPRSS2 and allows for subsequent escape from IFITM3. (A) Relative lentiviral pseudotype (PV) entry into 293T cells expressing ACE2-FLAG with or without co-expression of TMPRSS2. Entry into cells not transfected with TMPRSS2 normalised to 1. Statistics determined by multiple t-tests. ****, P ≤ 0.0001. (B-D) Relative PV entry into (D) 293T-ACE2, (E) Caco-2 or (F) Calu-3 cells pretreated with Amphotericin B (pink bars). Entry into untreated cells normalised to 1 (black bars). Statistics determined by multiple t-tests. *, 0.05 ≥ P > 0.01; **, 0.01 ≥ P > 0.001 (E) Relative PV entry into 293T cells overexpressing ACE2-FLAG and TMPRSS2, with or without IFITM3. Entry into cells not transfected with IFITM3 normalised to 1 (black bars). Statistics determined by multiple t-tests. **, 0.01 ≥ P > 0.001; ***, 0.001 ≥ P > 0.0001; ****, P ≤ 0.0001. (F) Replication kinetics of SARS-CoV-2 WT and ΔCS viruses in HAE cells. Cells were pretreated with control media or media containing amphotericin B for 1 hour then infected at an MOI of 0.1. Statistics were determined by one-way ANOVA with multiple comparisons on log transformed data. Black asterisks indicate statistical significance between no drug controls of WT and ΔCS while coloured asterisks indicate significance between no drug control or amphoB. DMSO controls were the same as from Figure 3D. *, 0.05 ≥ P > 0.01; **, 0.01 ≥ P > 0.001; ***, 0.001 ≥ P > 0.0001; ****, P ≤ 0.0001.

### The furin cleavage site of SARS-CoV-2 allows escape from IFITM2/3 proteins in TMPRSS2 expressing cells

The enhanced replication shown by SARS-CoV-2 with furin pre-cleaved spike to utilise TMPRSS2 to enter cells efficiently suggested this virus may prefer to enter cells via membranes near the cell surface or in the early endosome rather than be trafficked into late endosomes where it could be cleaved by host cathepsins. We hypothesised that the wild type virus may be avoiding host restriction factors in the endosome, such as IFITM2/3 proteins, which have previously been shown to be able to restrict SARS-CoV and SARS-CoV-2 entry (Huang et al., 2011; Zhao et al., 2020; Zheng et al., 2020). The antifungal agent amphotericin B (amphoB) has been well described as inhibiting the restriction imposed by IFITM proteins, particularly the endosomal/lysosomal localised IFITM2/3, potentially modulating the host membrane fluidity required for efficient restriction (Lin et al., 2013; Zhao et al., 2020; Zheng et al., 2020). We could also show that 293T-ACE2, Caco-2, Calu-3 and HAEs all constitutively expressed IFITM3, even in the absence of exogenous interferon (Figure 3E-H). We treated cells with amphoB and investigated the effect on PV entry. In 293T-ACE2 cells, entry of all coronavirus PV was improved by amphoB pre-treatment, with the largest effect exerted upon PVs with polybasic CS containing spikes (Figure 4B). Conversely, in Caco-2 and Calu-3 cells, the opposite effect was seen – entry of PVs with non-furin CS spikes was boosted, in some cases significantly by amphoB treatment, whereas there was little or no effect on the entry of PVs with furin CS containing spikes (Figure 4C,D).

Next, we took 293T cells and transiently co-transfected a combination of ACE2 and TMPRSS2 in the presence or absence of IFITM3. Entry of PVs with furin CS containing spikes were less inhibited by IFITM3 than those with spikes that could not be furin cleaved (Figure 4E, Supplementary Figure S1B).

Finally, we investigated the effect of amphoB treatment on SARS-CoV-2 replication in primary airway cells. AmphoB had no effect on WT virus replication, but greatly increased the replication of the ΔCS mutant (Figure 4F). This implies that IFITM proteins, such as IFITM3, are a major block for entry of viruses without furin CSs in these cells.

### The SARS-CoV-2 polybasic cleavage site promotes replication in the respiratory tract and transmission in a ferret model

To investigate whether the furin CS plays a role in the transmission of SARS-CoV-2, we used ferrets as an in vivo model. Ferrets are commonly used in transmission studies of respiratory pathogens such as influenza and, more recently, SARS-CoV-2 (Belser et al., 2018; Kim et al., 2020; Richard et al., 2020). Four ferrets per group were each infected intranasally with 10^5^ pfu of clonally purified stocks of the WT or ΔCS mutant SARS-CoV-2 viruses. After 24 hours, naïve contact ferrets were placed into each cage. All ferrets were nasal washed daily for the following 2 weeks and virus shedding in the nose was titrated by qRT-PCR and by TCID_50_ (Figure 5A,B, Supplementary Figure S2A,B). All eight directly inoculated ferrets shed virus robustly for 9-12 days (Figure 5A). The WT infected group shed more virus in the nose than ferrets infected with ΔCS virus, indicated by higher infectivity and higher E gene copy numbers, the latter significant at days 2-4. In the WT group, 2/4 contact ferrets became productively infected indicated by infectious virus, E gene loads in nasal wash, and seroconversion, whereas no transmission from donor ferret infected with ΔCS mutant virus was detected (Figure 5B, Supplementary Figures S2A-C). In nasal washes of the two remaining ferrets exposed to donors infected with WT virus, low E gene copy numbers were detected but no infectious virus was measured by TCID_50_, and these animals remained seronegative at 14 days post exposure, implying these ferrets were genuinely not infected in contrast to all directly infected animals and the two virus positive WT infected sentinels (Supplementary Figure S2C).

**Figure 5.**
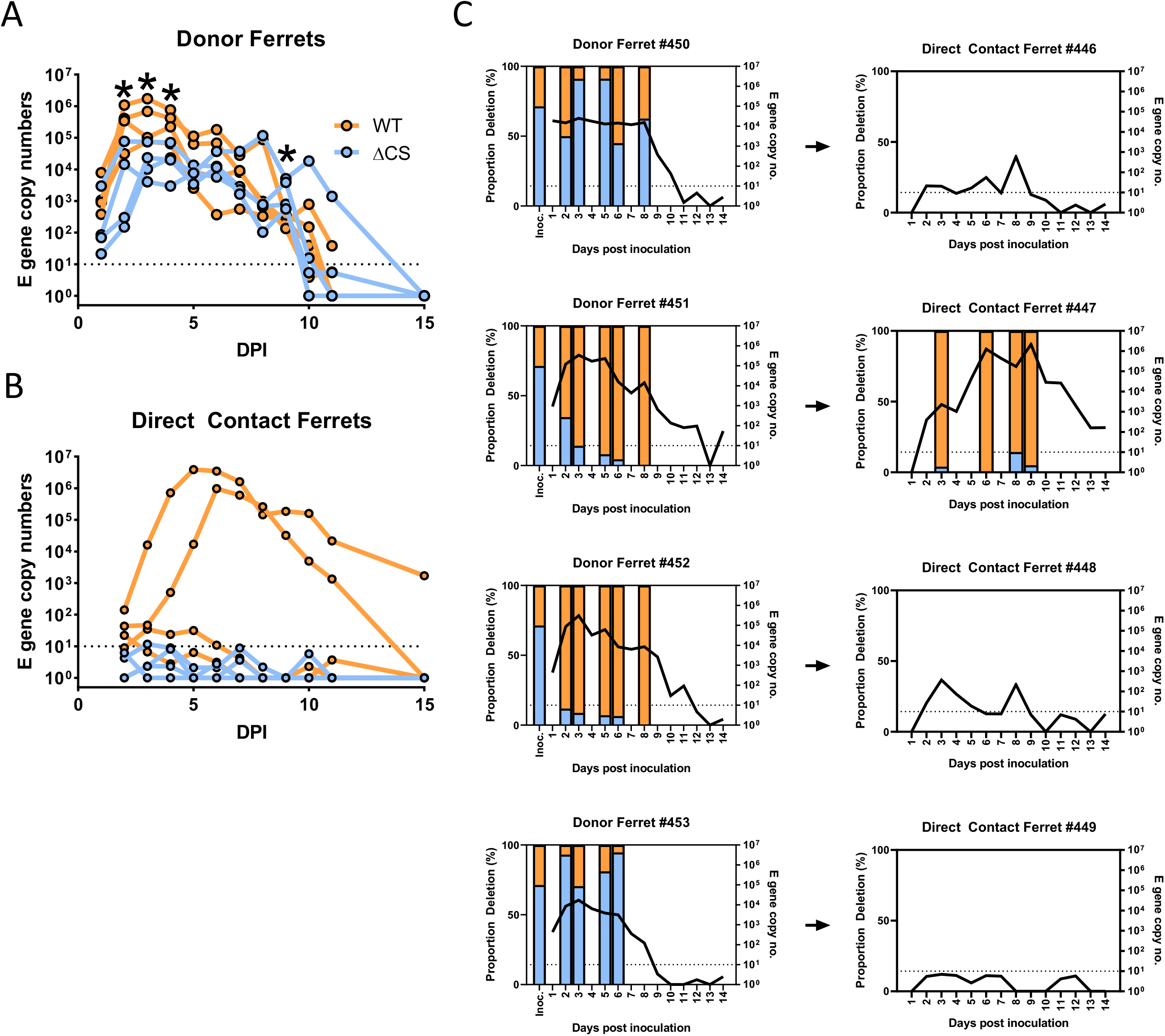
The furin cleavage site of SARS-CoV-2 allows for efficient replication and transmission in a ferret model. (A, B) Head to head transmission experiment of SARS-CoV-2 mix of WT and ΔCS in ferrets. In each group four individually housed donor ferrets were infected with X pfu of either WT or ΔCS SARS-CoV-2. One day post-inoculation naïve contact ferrets were added to each donor ferret. Ferrets were sampled by nasal wash daily and direct contact (A) and contact (B) ferret virus titres were determined by E gene qPCR. Statistics were determined by multiple t tests of the log transformed E gene copy numbers between each group. *, 0.05 ≥ P (C) Competition transmission experiment of SARS-CoV-2 mix of WT and ΔCS in ferrets. Four individually housed donor ferrets were infected with 105 pfu of virus mix containing ^~^70% ΔCS and ^~^30% WT. One day post-inoculation naïve contact ferrets were added to each donor ferret. Ferrets were sampled by nasal wash daily and virus titres were determined by E gene qPCR. For donors on day 2, 3, 5, 6 and 8, and for contacts on day 3, 6, 8 and 9 viral RNA across the S1/S2 cleavage site was deep sequenced. Where sequencing data was obtained bars showing the ratio of WT and deletion are shown.

Next, a competition assay was performed whereby four ferrets were inoculated intra-nasally with 10^5^ pfu of the previously described mixture of WT and ΔCS virus at a 30:70 ratio, as determined by deep sequencing. One day post-inoculation, naïve contact ferrets were cohoused with each directly inoculated animal and all animals were nasal washed daily. All directly inoculated ferrets became productively infected, shedding infectious virus and detectable E gene in nasal wash for between 9-12 days (Figure 5C). Interestingly, which virus genotype became dominant in the nasal washes of the directly infected ferrets appeared to vary stochastically; in two animals the WT virus became predominant by day 2, and levels of infectious virus and E gene detected in their nasal wash was highest. In the nasal wash of the other two directly inoculated animals, the ΔCS virus remained the majority species or outcompeted the WT over the course of the experiment. Productive transmission was only recorded in a single contact and interestingly, this animal was co-caged with one of the animals that was shedding predominantly WT virus. The ΔCS genotype was detectable in this single contact at low levels on day 3, 8 and 9, but at all times the WT genotype was clearly predominant. Furthermore, this was the only contact ferret to seroconvert, confirming the other 3 contact ferrets were not productively infected (Supplementary Figure S3D). No ferrets in either group showed appreciable fever or weight loss (Supplementary Figures S3A-C, S2D,E). Together these results suggest that the furin CS of SARS-CoV-2 is a determinant of transmission in the ferret model.

### SARS-CoV-2 spike variants with deletions or mutations in the polybasic cleavage site are detectable in human tissues

Finally, we wanted to investigate whether spike deletion mutants could be found in human clinical samples and, if so, whether they were more likely to be found in a particular organ. Initially we downloaded 100,000 genome sequences from GISAID and found 2 sequences from nasal swabs with CS deletions (Supplementary Table S1). Next, we sequenced the S1/S2 CS from 24 previously described post-mortem samples taken from five different post-mortems and including tissues from the respiratory and gastrointestinal tract, the brain, heart, bone marrow, kidney, tongue and spleen (Hanley et al., 2020). Sequencing revealed very low levels of viral RNA bearing different S1/S2 CS deletion (<1%) from heart and spleen tissue from separate patients (Supplementary Table S2). The three separate deletions reported in Supplementary Table S1 have not previously been reported but are similar to those seen upon passage in Vero E6 cells. OS5 deletes 4 amino acids after the CS similar to a deletion reported in a recent preprint (Sasaki et al., 2020); OS19-1 overlaps with most of ΔFlank and OS19-2 completely removes the S1/S2 site, similar to ΔCS. We have also observed identical deletions to OS19-2 upon passaging the clonal WT virus in Vero E6 cells. These results are consistent with the conclusion that S1/S2 cleavage site deletions can arise naturally in vivo, albeit at a very low rate.

## Discussion

An insertion of 4 amino acids in the SARS-CoV-2 spike protein created a suboptimal furin cleavage site (CS). Here, we propose a mechanism by which this conferred an advantage to the virus in the human airway and enabled efficient human-to-human transmission. We show that pre-cleavage of the spike during viral egress enhanced entry of progeny virions into TMPRSS2-expressing cells such as those abundant in respiratory tissue. TMPRSS2 cleaves spike at S2’ and facilitates entry at the cell surface (or possibly the early endosome) as opposed to entry through the endosome. This allows virus to avoid the otherwise potent IFITM restriction factors that reside in the late endosome and lysosome and inhibit viral membrane fusion in these compartments. Viruses that lack a polybasic S1/S2 CS cannot be cleaved by furin in producer cells and are thus forced to enter the next cell they infect through the endosome where the spike can be cleaved at S1/S2 and S2’ by cathepsins. However, pre-cleaved spike is not always advantageous: in cell types lacking TMPRSS2 expression, such as Vero E6, viruses without the furin CS gain an advantage, potentially because they are more stable, since spike cleavage may result in premature loss of the S1 subunit altogether and abrogate receptor binding. Our results show that, in contrast with WT SARS-CoV-2, a virus with a deleted furin CS did not replicate to high titres in the upper respiratory tract of ferrets and did not transmit to cohoused sentinel animals, in agreement with recent results from similar experiments with hamsters (Y. Zhu et al., 2020). It is not yet clear whether transmission is blocked due to the lower titres released from the directly inoculated donor ferrets, or to a lower ability to initiate infection in the TMPRSS2-rich cells of the nasal epithelium or a combination of these. We have found that furin CS deletions arise naturally in various different human organs during severe infection, but rarely and at low levels. Indeed, we note only 2 recorded genomes on GISAID out of 100,690 (as of 16/9/20) with furin CS deletions (Supplemental Table S1). Given the ease of loss of the furin CS in cell culture, the lack of these mutants in sequenced isolates is further evidence that the furin CS is essential for sustained transmission of SARS-CoV-2 in humans.

Presence of a furin CS at the S1/S2 junction is not uncommon in human coronaviruses; two of the four seasonal coronaviruses that transmit efficiently in humans, hCoV-HKU1 and hCoV-OC43, both contain furin CSs while MERS-CoV contains a suboptimal dibasic furin CS (Le Coupanec et al., 2015; Millet & Whittaker, 2014). However, the other two human seasonal coronaviruses, hCoV-229E and the ACE2 binding hCoV-NL63, do not contain furin CSs in their spike proteins seemingly without any detriment to transmissibility. Thus, furin-mediated cleavage of spike is not an absolute requirement for efficient human respiratory transmission. Furthermore, the original 2003 SARS-CoV lacked a furin CS and was clearly transmitted between people, although this usually occurred during a symptomatic phase rather than asymptomatically or pre-symptomatically, thus allowing the virus outbreak to be controlled by public health measures (Liu, Gayle, Wilder-Smith, & Rocklov, 2020). Several recent studies have investigated the pathogenicity of naturally occurring or engineered deletion mutants in the hamster model, and shown that viruses lacking the furin CS are attenuated for replication and pathogenicity (Johnson et al., 2020; Lau et al., 2020; Wang et al., 2020; Wong et al., 2020). We, like others, did not find that SARS-CoV-2 induced significant clinical signs in ferrets, so were unable to assess the effect of the CS mutation on viral pathogenesis here. Nonetheless, we did observe transmission between ferrets suggesting clinical signs are not required for transmission. A further single study also found a CS deletion mutant virus did not transmit between hamsters (Y. Zhu et al., 2020). Two studies suggest that the furin cleavage peptide motifs at the S1/S2 CS enables WT SARS-CoV-2 virus to utilise neuropilin as a cellular attachment/entry factor (Cantuti-Castelvetri et al., 2020; Daly et al., 2020). However, since 293T cells express neuropilin and lentiviral pseudotypes (PV) lacking the furin CS enter these cells more efficiently than PV containing it, this suggests that neuropilin is not responsible for the entry differences between furin cleavage mutants.

IFITM proteins are well characterised restriction factors of many enveloped viruses. Human IFITM3, for example, is a potent inhibitor of influenza infection and naturally occurring variants have been associated with severe influenza infection (Everitt et al., 2012; Huang et al., 2011; Zhang et al., 2013). IFITM proteins have also been shown to inhibit coronavirus entry (Huang et al., 2011; Wrensch, Winkler, & Pohlmann, 2014), although hCoV-OC43 and hCoV-HKU1 are unique in that IFITM proteins appear to act as entry factors to these viruses (Zhao et al., 2014; Zhao et al., 2018). There is growing evidence that IFITM proteins inhibit SARS-CoV-2 syncytia formation or pseudoviral or viral entry (Bozzo et al., 2020; Buchrieser et al., 2020; Shi et al., 2020), as we confirm here. This may partly explain the vulnerability of SARS-CoV-2 to IFN treatment (Mantlo, Bukreyeva, Maruyama, Paessler, & Huang, 2020). A recent preprint has suggested that under certain conditions IFITM proteins can have a modest pro-viral effect on SARS-CoV-2 virus growth (Bozzo et al., 2020). This is not inconsistent with our data which showed that amphotericin B mediated inhibition of IFITM proteins had no effect on, and if anything, slightly inhibited, WT SARS-CoV-2 virus replication in primary human cells.

Our study confirms TMPRSS2 as a potential drug target. Whilst inhibition of TMPRSS2 protease activity would not prevent infection via the endosome, using this pathway is detrimental to virus replication in airway cells. We have shown in this study that the protease inhibitor, camostat, is highly efficient at blocking SARS-CoV-2 replication in human airway cells and we note that clinical trials are ongoing [ClinicalTrials.gov Identifier: NCT04455815]. Our study also confirms the limitations of relying on Vero E6 cells as a system for developing classes of drugs such as entry inhibitors as they do not accurately reflect the preferred entry mechanism of SARS-CoV-2 into human airway cells (Hoffmann, Mosbauer, et al., 2020; T. Ou et al., 2020). Indeed the data here explains why chloroquine is ineffective in clinic against SARS-CoV-2 (Hoffmann, Mosbauer, et al., 2020), since during replication in the human airway WT SARS-CoV-2 has evolved to enter cells without the need for endosomal acidification. Monitoring wild coronaviruses will likely be important in predicting and preventing future pandemics. We suggest that a furin CS in the SARS lineage is a cause for concern. The polybasic insertion to the S1/S2 CS provides a significant fitness advantage in TMPRSS2 expressing cells and is likely essential for efficient human transmission.

## Materials and Methods

### Biosafety and ethics statement

All work performed was approved by the local genetic manipulation (GM) safety committee of Imperial College London, St. Mary’s Campus (centre number GM77), and the Health and Safety Executive of the United Kingdom, under reference CBA1.77.20.1. Animal research was carried out under a United Kingdom Home Office License, P48DAD9B4.

Human samples used in this research project were obtained from the Imperial College Healthcare Tissue Bank (ICHTB). ICHTB is supported by the NIHR Biomedical Research Centre based at Imperial College Healthcare NHS Trust and Imperial College London. ICHTB is approved by Wales REC3 to release human material for research (17/WA/0161), and the samples for this project (R20012) were issued from subcollection reference number MED_MO_20_011.

### Cells and viruses

African green monkey kidney cells (Vero E6) and human embryonic kidney cells (293T) were maintained in Dulbecco’s modified Eagle’s medium (DMEM), 10% fetal calf serum (FCS), 1% non-essential amino acids (NEAA), 1% penicillin-streptomycin (P/S). Human epithelial colorectal adenocarcarcinoma cells (Caco-2) and human lung cancer cells (Calu-3) were maintained in DMEM, 20% FCS, 1% NEAA, 1% P/S. VeroE6/TMPRSS2 cells were obtained from the Centre for AIDS Reagents (National Institute for Biological Standards and Control) (Matsuyama et al., 2020; Nao et al., 2019), and maintained in DMEM, 10% FCS, 1% NEAA, 1% P/S, 1 mg/ml Geneticin (G418). Air liquid interface Human airway epithelial cells (HAEs) were purchased from Epithelix and maintained in Mucilair cell culture medium (Epithelix). All cell lines were maintained at 37°C, 5% CO_2_

293T-hACE2 were generated by transducing 293Ts with an ACE2 expressing lentiviral vector, MT126, and selecting with 2 μg/ml puromycin, after selection cells were subsequently maintained with 1 μg/ml of puromycin.

The mixed SARS-CoV-2 WT/deletion virus mix was produced as previously described (Davidson et al., 2020). Briefly, the mix was generated by passaging the strain England/2/2020 (VE6-T), isolated by Public Health England (PHE), in Vero E6 cells whereby the deletion spontaneously arose (Davidson et al., 2020). The WT SARS-CoV-2 strain SARS-CoV-2 strain England/2/2020 (VE6-T) and the ΔCS mutant present in the original mixed stock were purified by serially diluting the stock (10-fold dilutions) in MEM supplemented with 2% FCS and adding the dilutions to either Vero E6 or Caco-2 cells in a 96 well plate. After 5 days incubation at 37 °C in 5% CO2, the culture supernatants in wells showing CPE at the highest dilution were again diluted and passaged on the same cells. After a further 5 days incubation, a 20 μl aliquot of culture supernatant from wells showing CPE at the highest dilution were used for RNA extraction and RT-PCR using a primer set designed to discriminate the WT and ΔCS mutant viruses. Culture supernatants containing either the WT or ΔCS mutant virus, with no sign of a mixed virus population were used to produce large scale stocks in Vero E6 cells. The presence of the expected virus in the stocks was verified by direct RNA sequencing using an Oxford Nanopore flow cell as previously described (Davidson et al., 2020). Clonally pure viruses were then further amplified by one additional passage in Vero E6/TMPRSS2 cells to make the working stocks of the viruses used throughout this study.

For plaque assays Vero E6 cells were used at 70-80% confluence. Cells were washed with PBS then serial dilutions of inoculum, diluted in serum-free DMEM, 1% NEAA, 1% P/S, were overlayed onto cells for one hour at 37°C. Inoculum was then removed and replaced with SARS-CoV-2 overlay media (1x minimal essential media [MEM], 0.2% w/v bovine serum albumin, 0.16% w/v NaHCO_3_, 10mM Hepes, 2mM L-Glutamine, 1x P/S, 0.6% w/v agarose). Plates were incubated for 3 days at 37°C before overlay was removed and cells were stained for 1 hour at RT in crystal violet solution.

To titrate virus by TCID_50_ Vero E6 cells were used at 70-80% confluence. Serial dilutions of virus, diluted in serum-free DMEM, 1% NEAA, 1% P/S, were added to each well and cells were left for 5 days before they were fixed with 2x crystal violet solution and analysed. 4 replicates of each sample were performed in tandem. TCID_50_ titres were determined by the Spearman-Kärbar method (Kärber, 1931).

### Plasmids and cloning

Lentiviral packaging constructs pCSLW and pCAGGs-GAGPOL were made as previously described (Long, Wright, Molesti, Temperton, & Barclay, 2015). The codon-optimised spike proteins of SARS-CoV-2, SARS-CoV and MERS-CoV were a kind gift from Professor Robin Shattock, Imperial College London (McKay et al., 2020). Mutant SARS-CoV-2 expression plasmids were generated by site-directed mutagenesis. The lentiviral expression vector for human ACE2, MT126, was a kind gift from Caroline Goujon, University of Montpellier. ACE was further cloned into pCAGGs with the addition of a C-terminal FLAG-tag. TMPRSS2 expression plasmid was a kind gift from Roger Reeves (Addgene plasmid #53887; http://n2t.net/addgene:53887; RRID:Addgene_53887,(Edie et al., 2018)).

### Syncytia formation assay

Vero E6 cells were seeded in 96-well plates (6.5×10^3^ cells per well) to reach 70-80% confluency on the subsequent day. Transfection was performed using 100 ng of expression plasmid using 0.3 μl of FuGENE HD Transfection Reagent (Promega E2311) in 20 μl of Opti-MEM medium (Life Technologies). At 48 hr after drug treatment, plates were washed in 100 μL/well of 1xPBS and fixed in 40 μl 4% PFA for 10 min at RT. After fixation cells were permeabilized in 0.1% Tritonx100 for 10 min at RT. Nuclei were stained using Hoechst 33342 (H3570 ThermoFisher), according to the manufacturer’s instructions.

Image acquisition was performed using the Operetta CLS high content screening microscope (Perkin Elmer) with a Zeiss 20x (NA=0.80) objective. A total of 25 fields per well were imaged for the Hoechst 33342 channel (Excitation (Ex) 365-385nm, Emission (Em) 430-500nm). Images were subsequently analysed, using the Harmony software (PerkinElmer). Images were first flatfield-corrected and nuclei were segmented using the “Find Nuclei” analysis module (Harmony). The thresholds for image segmentation were adjusted according to the signal-to-background ratio. Splitting coefficient was set in order to avoid splitting of overlapping nuclei (fused cells). All the cells that had a nuclear area greater than 3 times the average area of a single nucleus were considered as fused. Data were expressed as a percentage of fused cells by calculating the average number of fused cells normalized on the total number of cells per well.

### Lentiviral pseudotype assays

Lentiviral pseudotypes (PV) were generated as previously described (Long et al., 2015; Sumner et al., 2020). Briefly, 10cm^2^ dishes of 293T cells were co-transfected with a mixture of 1 μg of the HIV packaging plasmid pCAGGs-GAGPOL, 1.5 μg of the luciferase reporter construct, pCSLW and 1 μg of each envelope protein in pcDNA3.1. PV containing supernatants were harvested at 48- and 72-hours post-transfection, passed through a 0.45 μm filter, aliquoted and frozen at −80°C. When used, the pro-protein convertase inhibitor Decanoyl-RVKR-CMK (Calbiochem) was applied to cells at a concentration of 5 μM, 3 hours post-transfection. Concentrated PV were produced by ultracentrifugation at 100,000 x g for 2 hours over a 20% sucrose cushion.

Cells were transduced by PV for 48 hours before lysis with cell culture lysis buffer (Promega). Luciferase luminescence was read on a FLUOstar Omega plate reader (BMF Labtech) using the Luciferase Assay System (Promega). The cathepsin inhibitor E-64d (Sigma-Aldrich), the serine protease inhibitor camostat mesylate (Abcam), or the antifungal and IFITM3 inhibitor Amphotericin B (Sigma-Aldrich) was pre-applied to cells for 1 hour at a concentration of 50 μM before addition of PV.

### Deep sequencing using primer ID

RNA was extracted from ferret nasal washes or cell supernatants using the QIAamp Viral RNA Mini Kit (Qiagen) with carrier RNA. RNA was reverse transcribed using Superscript IV (Invitrogen) and a barcoded primer for Primer ID (TGCGTTGATACCACTGCTTTNNNNANNNNANNNNAACTGAATTTTCTGCACCAAG). Primer ID attaches a unique barcode to each cDNA molecule during reverse transcription and allows for PCR and sequencing error correction (Goldhill et al., 2019; Goldhill et al., 2018; Jabara, Jones, Roach, Anderson, & Swanstrom, 2011). PCR was performed using KOD polymerase (Merck) and the following primers (CAACTTACTCCTACTTGGCGT and XXXXTGCGTTGATACCACTGCTTT) giving a 272bp amplicon. XXXX was a 4-base barcode (CACA, GTTG, AGGA or TCTC) to allow for additional multiplexing. Samples were pooled and prepared for sequencing using NebNext Ultra II (NEB), then sequenced on an Illumina MiSeq with 300bp paired-end reads. Sequences were analysed in Geneious (v11) and a pipeline in R. Forward and reverse reads were paired using FLASh (https://ccb.jhu.edu/software/FLASH) before being mapped to a reference sequence and consensus sequences made for each barcode. A minimum cut-off of 3 reads per barcode was chosen. Raw sequences were deposited at www.ebi.ac.uk/ena, project number PRJEB40394. The analysis pipeline can be found at github.com/Flu1/Corona

### Deep sequencing from post-mortem samples

RNA from human post-mortem tissues from SARS-CoV-2 patients where COVID-19 was listed clinically as the cause of death were sourced and processed as previously described (Hanley et al., 2020). Briefly, fresh tissue was processed within biosafety level 3 facilities and total RNA was extracted using TRIzol (Invitrogen)-chloroform extraction followed by precipitation and purification using an RNeasy mini kit (Qiagen). RNA was reverse transcribed using Superscript IV (Invitrogen) and the following primer (GTCTTGGTCATAGACACTGGTAG). PCR was performed using KOD polymerase (Merck) and the following primers (GTCTTGGTCATAGACACTGGTAG and GGCTGTTTAATAGGGGCTGAAC) giving a 260bp amplicon. Samples were prepared for sequencing using NebNext Ultra II (NEB), then sequenced on an Illumina MiSeq with 300bp paired-end reads. Sequences were analysed in Geneious (v11) and a pipeline in R. Forward and reverse reads were paired using FLASh (https://ccb.jhu.edu/software/FLASH) before being mapped to a reference sequence. Raw sequences were deposited at www.ebi.ac.uk/ena, project number PRJEB40394. The analysis pipeline can be found at github.com/Flu1/Corona.

### Human Clinical Samples

A total of 100,690 SARS-CoV-2 genomes were downloaded from GISAID on 16/9/2020 and aligned to 234bp from the spike protein using Geneious. In frame deletions were identified in R and analysed to ensure that samples had not been passaged prior to sequencing. Code for this analysis can be found at github.com/Flu1/Corona.

### Ferret transmission studies

Ferret transmission studies were performed in a containment level 3 laboratory, using a bespoke isolator system (Bell Isolation Systems, U.K). Outbred female ferrets (16-20 weeks old), weighing 750-1000 g were used.

Prior to the study, ferrets were confirmed to be seronegative against SARS-CoV-2. Four donor ferrets were inoculated intranasally with 200 μl of 10^5^ pfu of virus mix while lightly anaesthetised with ketamine (22 mg/kg) and xylazine (0.9 mg/kg). To assess direct contact transmission one naïve direct contact ferrets were introduced into each cage 1-day post initial inoculation.

All animals were nasal washed daily, while conscious, by instilling 2 mL of PBS into the nostrils, the expectorate was collected into disposable 250 ml sample pots. Ferrets were weighed daily post-infection, and body temperature was measured daily via subcutaneous IPTT-300 transponder (Plexx B.V, Netherlands).

### Virus Neutralisation assay

The ability of ferret sera to neutralise wild type SARS-CoV-2 virus was assessed by neutralisation assay on Vero E6 cells. Heat-inactivated sera were serially diluted in assay diluent consisting of DMEM (Gibco, Thermo Fisher Scientific) with 1% penicillin-streptomycin (Thermo Fisher Scientific), 0.3% BSA fraction V (Thermo Fisher Scientific). Serum dilutions were incubated with 100 TCID_50_/well of virus in assay diluent for 1 h at RT and transferred to 96-well plates pre-seeded with Vero E6 cells. Serum dilutions were performed in duplicate. Plates were incubated at 37°C, 5% CO2 for 5 days before adding an equal volume of 2X crystal violet stain to wells for 1 h. Plates were washed, wells were scored for cytopathic effect and a neutralisation titre calculated as the reciprocal of the highest serum dilution at which full virus neutralisation occurred.

### qPCR

Viral RNA was extracted from Ferret nasal washes using the Qiagen Viral RNA mini kit, according to manufacturer’s instructions.

Quantitative real-time RT-PCR (qRT-PCR) was performed using 7500 Real Time PCR system (ABI) in 20 μl reactions using AgPath-ID One-Step RT-PCR Reagents 10 μl RT-PCR buffer (2X) (Thermo Fisher), 5μl of RNA, 1 μl forward (5' ACAGGTACGTTAATAGTTAATAGCGT 3') and reverse primers (5' ATATTGCAGCAGTACGCACACA 3') and 0.5 μl probe (5’ FAM-ACACTAGCCATCCTTACTGCGCTTCG -BBQ 3’). The following conditions were used: 45°C for 10 min, 1 cycle; 95°C for 15 min, 1 cycle; 95°C for 15 sec then 58°C for 30 sec, 45 cycles. For each sample, the C_t_ value for the target E gene was determined. Based on the standard curves, absolute E gene copy numbers were calculated.

RNA was extracted from cells using RNA extraction kits (QIAGEN, RNeasy Mini Kit, cat. 74106) following the manufacturer’s instructions. Complementary DNA (cDNA) was synthesized in a reverse transcription step using Oligo-dT (RevertAid First Strand cDNA Synthesis, ThermoScientific, cat: K1621). To quantify mRNA levels, real-time quantitative PCR analysis with a gene specific primer pair using SYBR green PCR mix (Applied Biosystems, cat: 4385612) was performed and data was analysed on the Applied Biosystems ViiATM 7 Real-Time PCR System. The following primers were used:

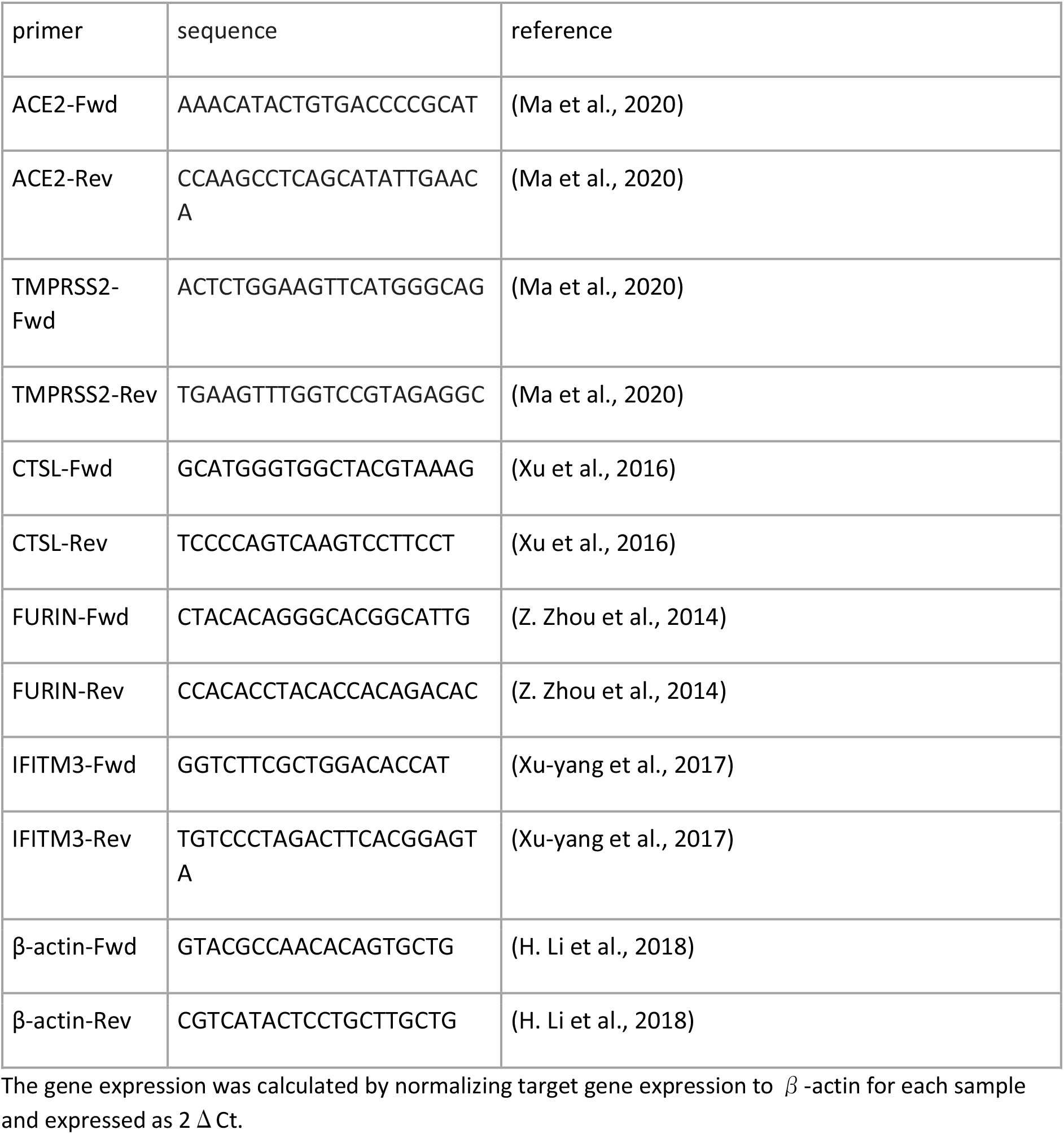

### Western Blotting

To investigate cleavage of spike protein 293T cells transfected with 2.5 μg of spike expression plasmids (or empty vector). After 48 hours cells were lysed in RIPA buffer (150mM NaCl, 1% NP-40, 0.5% sodium deoxycholate, 0.1% SDS, 50mM TRIS, pH 7.4) supplemented with an EDTA-free protease inhibitor cocktail tablet (Roche). Cell lysates were then mixed with 4x Laemmli sample buffer (Bio-Rad) with 10% β-mercaptoethanol. Concentrated PV as described above were also diluted in Laemmli buffer.

Membranes were probed with mouse anti-FLAG (F1804, Sigma), mouse anti-tubulin (abcam; ab7291), mouse anti-p24 (abcam; ab9071), rabbit anti-TMPRSS2 (abcam; ab92323), rabbit anti-Fragilis/IFITM3 (abcam; ab109429), rabbit anti-SARS spike protein (NOVUS; NB100-56578) or rabbit anti-SARS-CoV-2 nucleocapsid (SinoBiological; 40143-R019). Near infra-red (NIR) secondary antibodies, IRDye^®^ 680RD Goat anti-mouse (abcam; ab216776), IRDye^®^ 680RD Goat anti-rabbit (abcam; ab216777), IRDye^®^ 800CW Goat anti-mouse (abcam; ab216772), or IRDye^®^ 800CW Goat anti-rabbit (abcam; ab216773)) were subsequently used. Western blots were visualised using an Odyssey Imaging System (LI-COR Biosciences).

## Acknowledgments

SARS-CoV-2 virus was initially provided by Public Health England and we would like to thank Maria Zambon, Robin Gopal and Monika Patel for their help. This work was supported by BBSRC grants BB/R013071/1 (TPP, WB); BB/R007292/1 (LB, WB); BB/S008292/1 (JB, WB); BB/M02542X/1 (ADD, DAM) and Wellcome Trust grants 205100 (DHG, RYSD, WB); 200187 (JZ, RF, WB). This work was also supported by MRC grant MR/R020566/1 (MKW, ADD) and US FDA grant HHSF223201510104C (ADD, DAM). Additional support was provided from a grant from the King’s College London King’s Together Programme and the King’s College London BHF Centre of Research Excellence grant RE/18/2/34213 to MG. OCS was supported by a Wellcome Trust studentship, RK was supported by Wellcome fellowship 216353/Z/19/Z, RP was supported by an MRC DTP studentship, JAH was supported by a BBSRC DTP studentship and ES was supported by an Imperial College President’s Scholarship.

## Supplementary Figure Legends

**Supplementary Figure S1.**
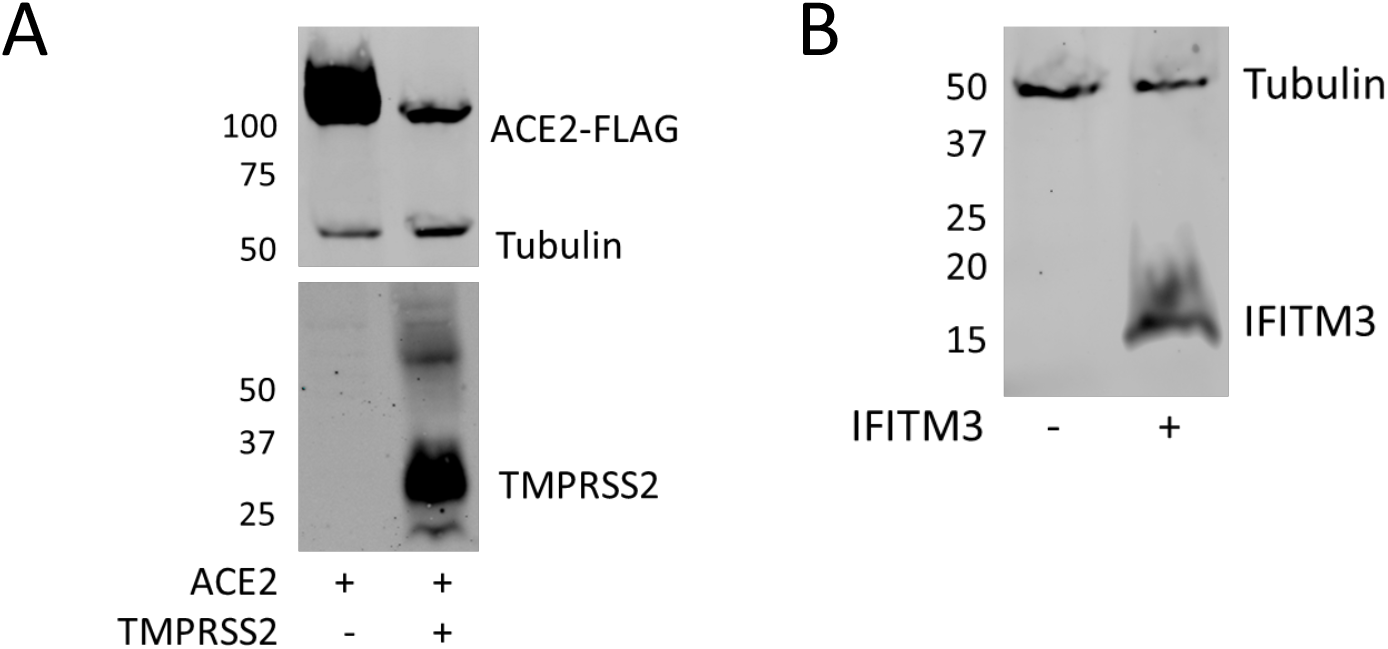
Expression of transfected ACE2, TMPRSS2 and IFITM3. (A) Expression of TMPRSS2 and ACE2-FLAG in 293T cells shown by western blot (from Figure 4A). (B) Expression of IFITM3 in 293T cells shown by western blot (from Figure 4G)

**Supplementary Figure S2.**
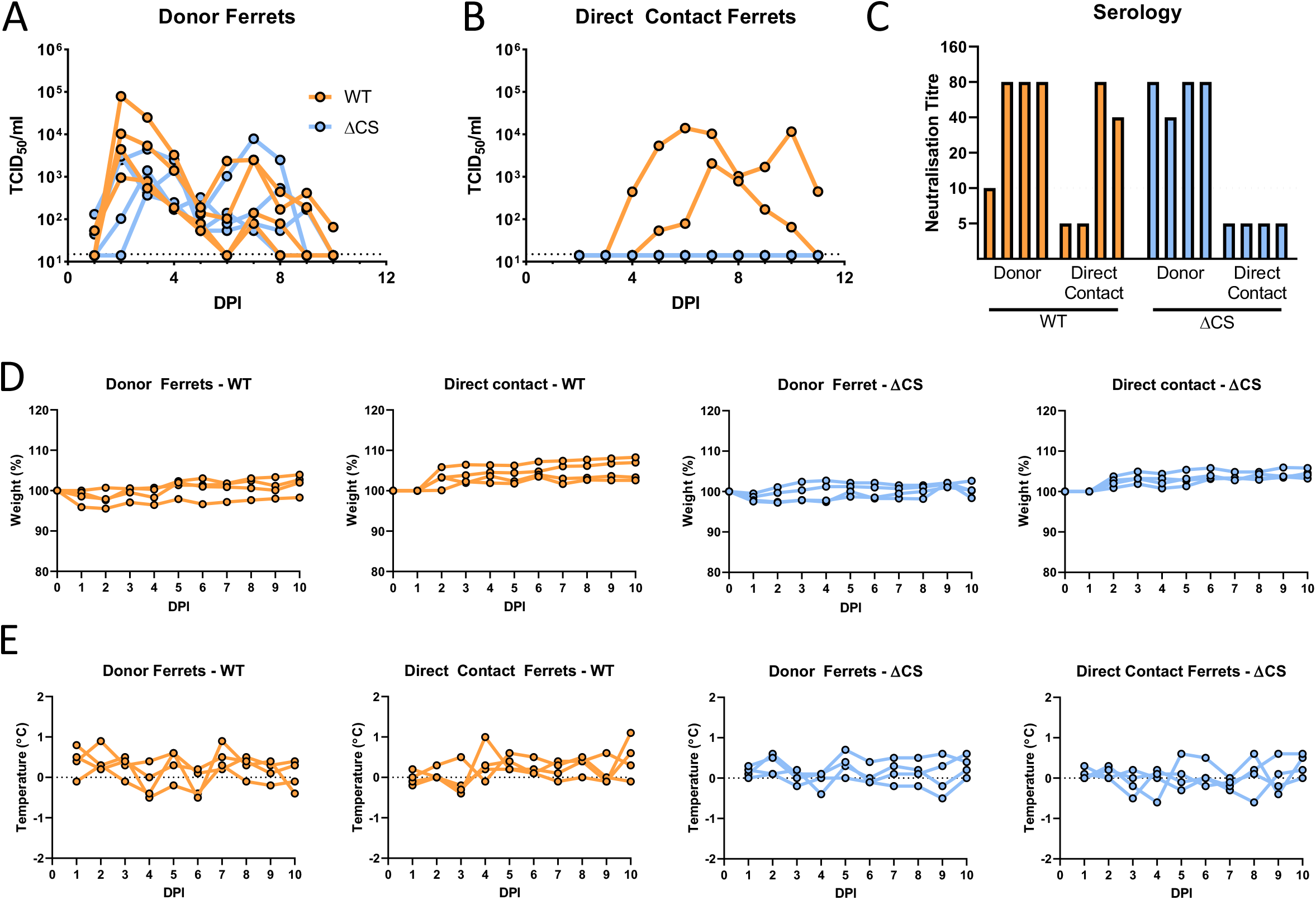
Ferret head-to-head transmission experiment addition shedding data and clinical data. (A, B) Infectious virus titres of WT (orange) or ΔCS (blue) taken from donor ferret (A) and direct contact ferret (B) nasal washes as determined by TCID_50_. Dotted line indicates limit of detection of infectious virus. (C) Microneutralisation assay showing ferret post-infection serology. Threshold of detection was a neutralisation titre of 10 (dotted line). Serum taken 14 days post-infection. (D) Changes in donor and direct contact ferret body weights during the duration of the infection. (E) Changes in donor and direct contact ferret body temperatures during the duration of the infection.

**Supplementary Figure S3.**
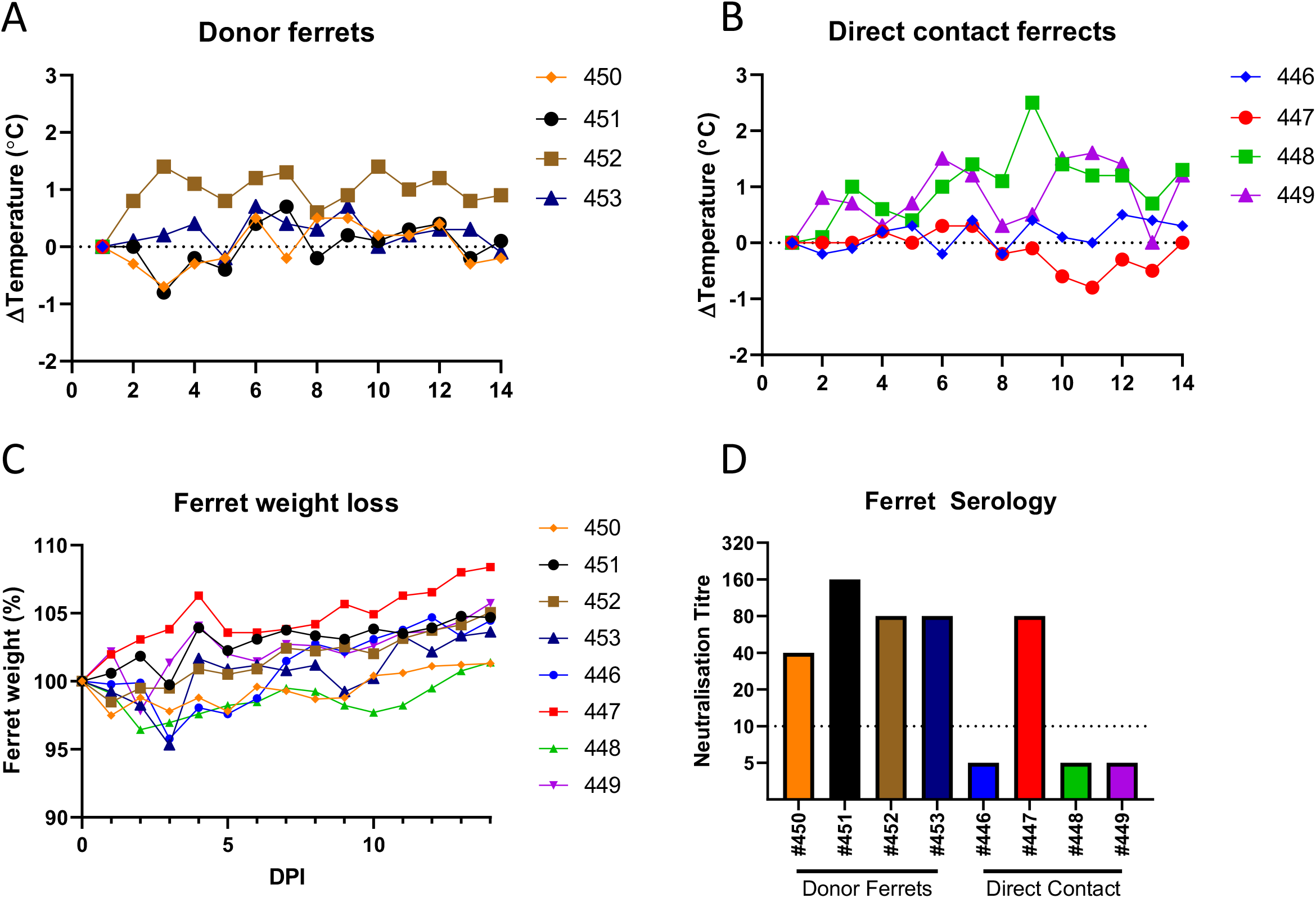
Ferret competition transmission experiment clinical data. (A, B) Changes in donor (A) and direct contact (B) ferret body temperatures during the duration of the infection. (C) Changes in ferret body weight (in percentage) during the duration of the infection. (D) Microneutralisation assay showing ferret post-infection serology. Threshold of detection was a neutralisation titre of 10 (dotted line). Serum taken 14 days post-infection.

**Supplementary Table 1.**
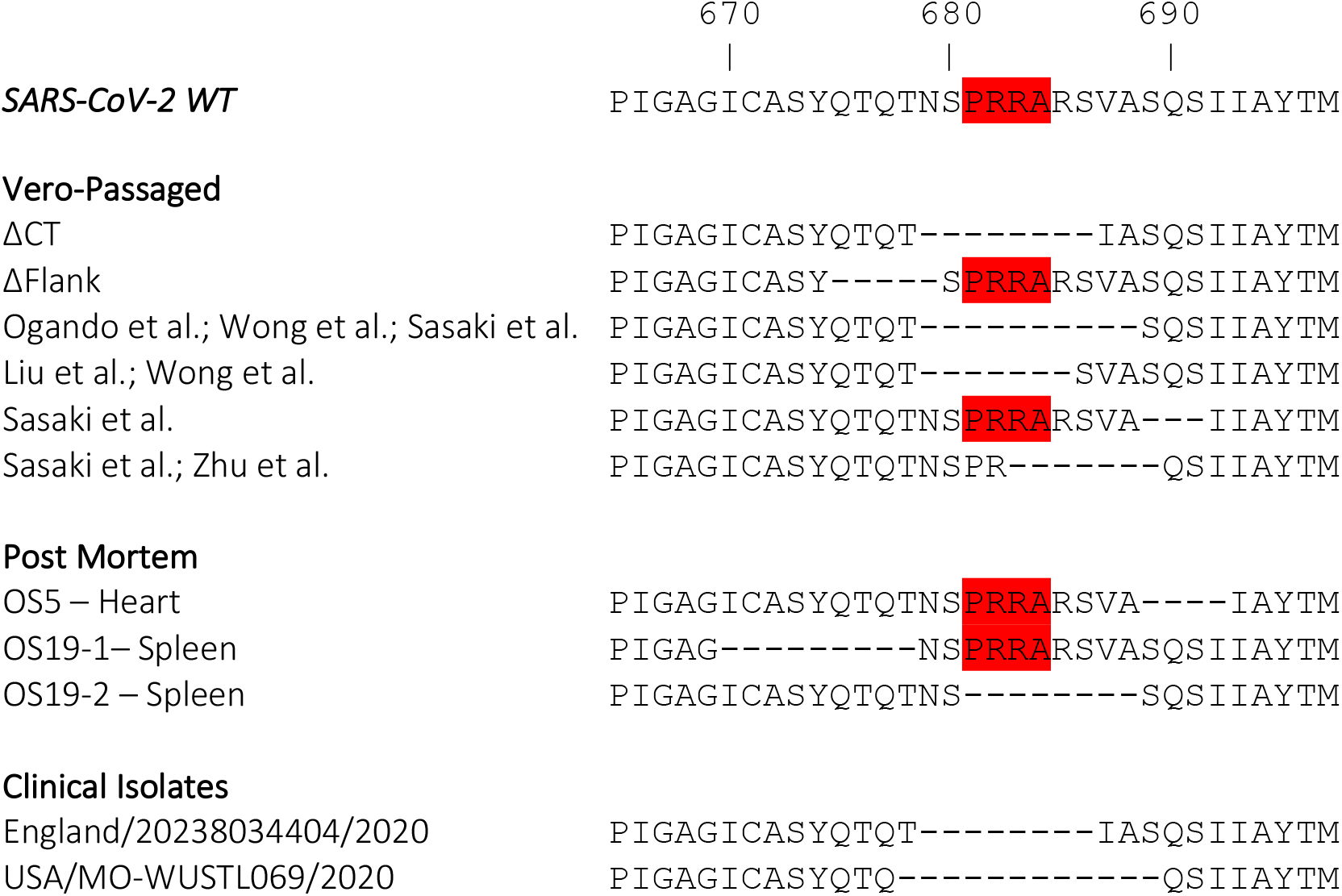
S1/S2 cleavage site deletions reported following viral passage in cell culture or from clinical samples.

**Supplementary Table 2.** Post-mortem samples sequenced for S1/S2 cleavage site deletions.

| Sample type | Post mortem case | Ct value by E gene qRT-PCR | Deletions detected? |
| --- | --- | --- | --- |
| bone marrow | PM3 | 30.2 | N |
| brain | PM2 | 25.4 | N |
| heart | PM1 | 18.8 | N |
|  | PM2 | 21.2 | Y - OS5 |
|  | PM6 | 17.7 | N |
| ileum | PM1 | 31.3 | N |
|  | PM2 | 28.3 | N |
|  | PM4 | 31.7 | N |
|  | PM6 | 25.1 | N |
| kidney | PM4 | 31.4 | N |
|  | PM6 | 25.8 | N |
| lung | PM1 | 30.2 | N |
|  | PM3 | 19.7 | N |
|  | PM6 | 11.5 | N |
| nasal | PM1 | 32.1 | N |
|  | PM3 | 30.8 | N |
|  | PM4 | 23.8 | N |
| spleen | PM3 | 28.5 | Y - OS19-1, OS19-2 |
| tongue | PM3 | 24.3 | N |
|  | PM4 | 24.3 | N |
|  | PM6 | 21.9 | N |
| trachea | PM1 | 32.7 | N |
|  | PM3 | 19.9 | N |
|  | PM4 | 23.5 | N |

## References

Andersen, K. G., Rambaut, A., Lipkin, W. I., Holmes, E. C., & Garry, R. F. (2020). The proximal origin of SARS-CoV-2. Nat Med, 26(4), 450–452. doi:10.1038/s41591-020-0820-9

Belouzard, S., Chu, V. C., & Whittaker, G. R. (2009). Activation of the SARS coronavirus spike protein via sequential proteolytic cleavage at two distinct sites. Proc Natl Acad Sci U S A, 106(14), 5871–5876. doi:10.1073/pnas.0809524106

Belser, J. A., Barclay, W., Barr, I., Fouchier, R. A. M., Matsuyama, R., Nishiura, H., … Yen, H. L. (2018). Ferrets as Models for Influenza Virus Transmission Studies and Pandemic Risk Assessments. Emerg Infect Dis, 24(6), 965–971. doi:10.3201/eid2406.172114

Bertram, S., Glowacka, I., Blazejewska, P., Soilleux, E., Allen, P., Danisch, S., … Pohlmann, S. (2010). TMPRSS2 and TMPRSS4 facilitate trypsin-independent spread of influenza virus in Caco-2 cells. J Virol, 84(19), 10016–10025. doi:10.1128/JVI.00239-10

Boni, M. F., Lemey, P., Jiang, X., Lam, T. T., Perry, B. W., Castoe, T. A., … Robertson, D. L. (2020). Evolutionary origins of the SARS-CoV-2 sarbecovirus lineage responsible for the COVID-19 pandemic. Nat Microbiol. doi:10.1038/s41564-020-0771-4

Bozzo, C. P., Nchioua, R., Volcic, M., Wettstein, L., Weil, T., Krüger, J., … Kirchhoff, F. (2020). IFITM proteins promote SARS-CoV-2 infection of human lung cells. bioRxiv, 2020.2008.2018.255935. doi:10.1101/2020.08.18.255935

Buchrieser, J., Dufloo, J., Hubert, M., Monel, B., Planas, D., Rajah, M. M., … Schwartz, O. (2020). Syncytia formation by SARS-CoV-2 infected cells. bioRxiv, 2020.2007.2014.202028. doi:10.1101/2020.07.14.202028

Cantuti-Castelvetri, L., Ojha, R., Pedro, L. D., Djannatian, M., Franz, J., Kuivanen, S., … Simons, M. (2020). Neuropilin-1 facilitates SARS-CoV-2 cell entry and provides a possible pathway into the central nervous system. bioRxiv, 2020.2006.2007.137802. doi:10.1101/2020.06.07.137802

Coutard, B., Valle, C., de Lamballerie, X., Canard, B., Seidah, N. G., & Decroly, E. (2020). The spike glycoprotein of the new coronavirus 2019-nCoV contains a furin-like cleavage site absent in CoV of the same clade. Antiviral Res, 176, 104742. doi:10.1016/j.antiviral.2020.104742

Daly, J. L., Simonetti, B., Antón-Plágaro, C., Kavanagh Williamson, M., Shoemark, D. K., Simón-Gracia, L., … Yamauchi, Y. (2020). Neuropilin-1 is a host factor for SARS-CoV-2 infection. bioRxiv, 2020.2006.2005.134114. doi:10.1101/2020.06.05.134114

Davidson, A. D., Williamson, M. K., Lewis, S., Shoemark, D., Carroll, M. W., Heesom, K. J., … Matthews, D. A. (2020). Characterisation of the transcriptome and proteome of SARS-CoV-2 reveals a cell passage induced in-frame deletion of the furin-like cleavage site from the spike glycoprotein. Genome Med, 12(1), 68. doi:10.1186/s13073-020-00763-0

Edie, S., Zaghloul, N. A., Leitch, C. C., Klinedinst, D. K., Lebron, J., Thole, J. F., … Reeves, R. H. (2018). Survey of Human Chromosome 21 Gene Expression Effects on Early Development in Danio rerio. G3 (Bethesda), 8(7), 2215–2223. doi:10.1534/g3.118.200144

Everitt, A. R., Clare, S., Pertel, T., John, S. P., Wash, R. S., Smith, S. E., … Kellam, P. (2012). IFITM3 restricts the morbidity and mortality associated with influenza. Nature, 484(7395), 519–523. doi:10.1038/nature10921

Goldhill, D. H., Langat, P., Xie, H., Galiano, M., Miah, S., Kellam, P., … Barclay, W. S. (2019). Determining the Mutation Bias of Favipiravir in Influenza Virus Using Next-Generation Sequencing. J Virol, 93(2). doi:10.1128/JVI.01217-18

Goldhill, D. H., Te Velthuis, A. J. W., Fletcher, R. A., Langat, P., Zambon, M., Lackenby, A., & Barclay, W. S. (2018). The mechanism of resistance to favipiravir in influenza. Proc Natl Acad Sci U S A, 115(45), 11613–11618. doi:10.1073/pnas.1811345115

Hanley, B., Naresh, K. N., Roufosse, C., Nicholson, A. G., Weir, J., Cooke, G. S., … Osborn, M. (2020). Histopathological findings and viral tropism in UK patients with severe fatal COVID-19: a post-mortem study. Lancet Microbe. doi:10.1016/S2666-5247(20)30115-4

Hoffmann, M., Kleine-Weber, H., & Pohlmann, S. (2020). A Multibasic Cleavage Site in the Spike Protein of SARS-CoV-2 Is Essential for Infection of Human Lung Cells. Mol Cell. doi:10.1016/j.molcel.2020.04.022

Hoffmann, M., Kleine-Weber, H., Schroeder, S., Kruger, N., Herrler, T., Erichsen, S., … Pohlmann, S. (2020). SARS-CoV-2 Cell Entry Depends on ACE2 and TMPRSS2 and Is Blocked by a Clinically Proven Protease Inhibitor. Cell, 181(2), 271–280 e278. doi:10.1016/j.cell.2020.02.052

Hoffmann, M., Mosbauer, K., Hofmann-Winkler, H., Kaul, A., Kleine-Weber, H., Kruger, N., … Pohlmann, S. (2020). Chloroquine does not inhibit infection of human lung cells with SARS-CoV-2. Nature. doi:10.1038/s41586-020-2575-3

Holshue, M. L., DeBolt, C., Lindquist, S., Lofy, K. H., Wiesman, J., Bruce, H., … Washington State -nCo, V. C. I. T. (2020). First Case of 2019 Novel Coronavirus in the United States. N Engl J Med, 382(10), 929–936. doi:10.1056/NEJMoa2001191

Huang, I. C., Bailey, C. C., Weyer, J. L., Radoshitzky, S. R., Becker, M. M., Chiang, J. J., … Farzan, M. (2011). Distinct patterns of IFITM-mediated restriction of filoviruses, SARS coronavirus, and influenza A virus. PLoS pathogens, 7(1), e1001258. doi:10.1371/journal.ppat.1001258

Jabara, C. B., Jones, C. D., Roach, J., Anderson, J. A., & Swanstrom, R. (2011). Accurate sampling and deep sequencing of the HIV-1 protease gene using a Primer ID. Proc Natl Acad Sci U S A, 108(50), 20166–20171. doi:10.1073/pnas.1110064108

Johnson, B. A., Xie, X., Kalveram, B., Lokugamage, K. G., Muruato, A., Zou, J., … Menachery, V. D. (2020). Furin Cleavage Site Is Key to SARS-CoV-2 Pathogenesis. bioRxiv, 2020.2008.2026.268854. doi:10.1101/2020.08.26.268854

Kärber, G. (1931). Beitrag zur kollektiven Behandlung pharmakologischer Reihenversuche. Naunyn-Schmiedebergs Archiv für experimentelle Pathologie und Pharmakologie, 162(4), 480–483. doi:10.1007/BF01863914

Kim, Y. I., Kim, S. G., Kim, S. M., Kim, E. H., Park, S. J., Yu, K. M., … Choi, Y. K. (2020). Infection and Rapid Transmission of SARS-CoV-2 in Ferrets. Cell Host Microbe, 27(5), 704–709 e702. doi:10.1016/j.chom.2020.03.023

Klimstra, W. B., Tilston-Lunel, N. L., Nambulli, S., Boslett, J., McMillen, C. M., Gilliland, T., … Duprex, W. P. (2020). SARS-CoV-2 growth, furin-cleavage-site adaptation and neutralization using serum from acutely infected hospitalized COVID-19 patients. J Gen Virol. doi:10.1099/jgv.0.001481

Lau, S. Y., Wang, P., Mok, B. W., Zhang, A. J., Chu, H., Lee, A. C., … Chen, H. (2020). Attenuated SARS-CoV-2 variants with deletions at the S1/S2 junction. Emerg Microbes Infect, 9(1), 837–842. doi:10.1080/22221751.2020.1756700

Le Coupanec, A., Desforges, M., Meessen-Pinard, M., Dube, M., Day, R., Seidah, N. G., & Talbot, P. J. (2015). Cleavage of a Neuroinvasive Human Respiratory Virus Spike Glycoprotein by Proprotein Convertases Modulates Neurovirulence and Virus Spread within the Central Nervous System. PLoS pathogens, 11(11), e1005261. doi:10.1371/journal.ppat.1005261

Li, H., Bradley, K. C., Long, J. S., Frise, R., Ashcroft, J. W., Hartgroves, L. C., … Barclay, W. S. (2018). Internal genes of a highly pathogenic H5N1 influenza virus determine high viral replication in myeloid cells and severe outcome of infection in mice. PLoS pathogens, 14(1), e1006821. doi:10.1371/journal.ppat.1006821

Li, W., Moore, M. J., Vasilieva, N., Sui, J., Wong, S. K., Berne, M. A., … Farzan, M. (2003). Angiotensin-converting enzyme 2 is a functional receptor for the SARS coronavirus. Nature, 426(6965), 450–454. doi:10.1038/nature02145

Lin, T. Y., Chin, C. R., Everitt, A. R., Clare, S., Perreira, J. M., Savidis, G., … Brass, A. L. (2013). Amphotericin B increases influenza A virus infection by preventing IFITM3-mediated restriction. Cell Rep, 5(4), 895–908. doi:10.1016/j.celrep.2013.10.033

Liu, Y., Gayle, A. A., Wilder-Smith, A., & Rocklov, J. (2020). The reproductive number of COVID-19 is higher compared to SARS coronavirus. J Travel Med, 27(2). doi:10.1093/jtm/taaa021

Liu, Z., Zheng, H., Lin, H., Li, M., Yuan, R., Peng, J., … Lu, J. (2020). Identification of common deletions in the spike protein of SARS-CoV-2. J Virol. doi:10.1128/JVI.00790-20

Long, J., Wright, E., Molesti, E., Temperton, N., & Barclay, W. (2015). Antiviral therapies against Ebola and other emerging viral diseases using existing medicines that block virus entry. F1000Res, 4, 30. doi:10.12688/f1000research.6085.2

Ma, D., Chen, C. B., Jhanji, V., Xu, C., Yuan, X. L., Liang, J. J., … Ng, T. K. (2020). Expression of SARS-CoV-2 receptor ACE2 and TMPRSS2 in human primary conjunctival and pterygium cell lines and in mouse cornea. Eye (Lond), 34(7), 1212–1219. doi:10.1038/s41433-020-0939-4

Mantlo, E., Bukreyeva, N., Maruyama, J., Paessler, S., & Huang, C. (2020). Antiviral activities of type I interferons to SARS-CoV-2 infection. Antiviral Res, 179, 104811. doi:10.1016/j.antiviral.2020.104811

Matsuyama, S., Nagata, N., Shirato, K., Kawase, M., Takeda, M., & Taguchi, F. (2010). Efficient activation of the severe acute respiratory syndrome coronavirus spike protein by the transmembrane protease TMPRSS2. J Virol, 84(24), 12658–12664. doi:10.1128/JVI.01542-10

Matsuyama, S., Nao, N., Shirato, K., Kawase, M., Saito, S., Takayama, I., … Takeda, M. (2020). Enhanced isolation of SARS-CoV-2 by TMPRSS2-expressing cells. Proc Natl Acad Sci U S A, 117(13), 7001–7003. doi:10.1073/pnas.2002589117

McKay, P. F., Hu, K., Blakney, A. K., Samnuan, K., Brown, J. C., Penn, R., … Shattock, R. J. (2020). Self-amplifying RNA SARS-CoV-2 lipid nanoparticle vaccine candidate induces high neutralizing antibody titers in mice. Nat Commun, 11(1), 3523. doi:10.1038/s41467-020-17409-9

Millet, J. K., & Whittaker, G. R. (2014). Host cell entry of Middle East respiratory syndrome coronavirus after two-step, furin-mediated activation of the spike protein. Proc Natl Acad Sci U S A, 111(42), 15214–15219. doi:10.1073/pnas.1407087111

Nao, N., Sato, K., Yamagishi, J., Tahara, M., Nakatsu, Y., Seki, F., … Takeda, M. (2019). Consensus and variations in cell line specificity among human metapneumovirus strains. PLoS One, 14(4), e0215822. doi:10.1371/journal.pone.0215822

Ogando, N. S., Dalebout, T. J., Zevenhoven-Dobbe, J. C., Limpens, R., van der Meer, Y., Caly, L., … Snijder, E. J. (2020). SARS-coronavirus-2 replication in Vero E6 cells: replication kinetics, rapid adaptation and cytopathology. J Gen Virol. doi:10.1099/jgv.0.001453

Ou, T., Mou, H., Zhang, L., Ojha, A., Choe, H., & Farzan, M. (2020). Hydroxychloroquine-mediated inhibition of SARS-CoV-2 entry is attenuated by TMPRSS2. bioRxiv, 2020.2007.2022.216150. doi:10.1101/2020.07.22.216150

Ou, X., Liu, Y., Lei, X., Li, P., Mi, D., Ren, L., … Qian, Z. (2020). Characterization of spike glycoprotein of SARS-CoV-2 on virus entry and its immune cross-reactivity with SARS-CoV. Nat Commun, 11(1), 1620. doi:10.1038/s41467-020-15562-9

Park, J. E., Li, K., Barlan, A., Fehr, A. R., Perlman, S., McCray, P. B., Jr., & Gallagher, T. (2016). Proteolytic processing of Middle East respiratory syndrome coronavirus spikes expands virus tropism. Proc Natl Acad Sci U S A, 113(43), 12262–12267. doi:10.1073/pnas.1608147113

Richard, M., Kok, A., de Meulder, D., Bestebroer, T. M., Lamers, M. M., Okba, N. M. A., … Herfst, S. (2020). SARS-CoV-2 is transmitted via contact and via the air between ferrets. Nat Commun, 11(1), 3496. doi:10.1038/s41467-020-17367-2

Sasaki, M., Uemura, K., Sato, A., Toba, S., Sanaki, T., Maenaka, K., … Sawa, H. (2020). SARS-CoV-2 variants with mutations at the S1/S2 cleavage site are generated *in vitro* during propagation in TMPRSS2-deficient cells. bioRxiv, 2020.2008.2028.271163. doi:10.1101/2020.08.28.271163

Shang, J., Wan, Y., Luo, C., Ye, G., Geng, Q., Auerbach, A., & Li, F. (2020). Cell entry mechanisms of SARS-CoV-2. Proc Natl Acad Sci U S A. doi:10.1073/pnas.2003138117

Shi, G., Kenney, A. D., Kudryashova, E., Zhang, L., Hall-Stoodley, L., Robinson, R. T., … Yount, J. S. (2020). Opposing activities of IFITM proteins in SARS-CoV-2 infection. bioRxiv, 2020.2008.2011.246678. doi:10.1101/2020.08.11.246678

Shirato, K., Kawase, M., & Matsuyama, S. (2013). Middle East respiratory syndrome coronavirus infection mediated by the transmembrane serine protease TMPRSS2. J Virol, 87(23), 12552–12561. doi:10.1128/JVI.01890-13

Shulla, A., Heald-Sargent, T., Subramanya, G., Zhao, J., Perlman, S., & Gallagher, T. (2011). A transmembrane serine protease is linked to the severe acute respiratory syndrome coronavirus receptor and activates virus entry. J Virol, 85(2), 873–882. doi:10.1128/JVI.02062-10

Simmons, G., Gosalia, D. N., Rennekamp, A. J., Reeves, J. D., Diamond, S. L., & Bates, P. (2005). Inhibitors of cathepsin L prevent severe acute respiratory syndrome coronavirus entry. Proc Natl Acad Sci U S A, 102(33), 11876–11881. doi:10.1073/pnas.0505577102

Sumner, R. P., Harrison, L., Touizer, E., Peacock, T. P., Spencer, M., Zuliani-Alvarez, L., & Towers, G. J. (2020). Disrupting HIV-1 capsid formation causes cGAS sensing of viral DNA. EMBO J, e103958. doi:10.15252/embj.2019103958

Wang, P., Lau, S.-Y., Deng, S., Chen, P., Mok, B. W.-Y., Zhang, A. J., … Chen, H. (2020). Pathogenicity, immunogenicity, and protective ability of an attenuated SARS-CoV-2 variant with a deletion at the S1/S2 junction of the spike protein. bioRxiv, 2020.2008.2024.264192. doi:10.1101/2020.08.24.264192

Wong, Y. C., Lau, S. Y., Wang To, K. K., Mok, B. W. Y., Li, X., Wang, P., … Chen, Z. (2020). Natural transmission of bat-like SARS-CoV-2PRRA variants in COVID-19 patients. Clin Infect Dis. doi:10.1093/cid/ciaa953

Wrensch, F., Winkler, M., & Pohlmann, S. (2014). IFITM proteins inhibit entry driven by the MERS-coronavirus spike protein: evidence for cholesterol-independent mechanisms. Viruses, 6(9), 3683–3698. doi:10.3390/v6093683

Xia, S., Lan, Q., Su, S., Wang, X., Xu, W., Liu, Z., … Jiang, S. (2020). The role of furin cleavage site in SARS-CoV-2 spike protein-mediated membrane fusion in the presence or absence of trypsin. Signal Transduct Target Ther, 5(1), 92. doi:10.1038/s41392-020-0184-0

Xu-yang, Z., Pei-yu, B., Chuan-tao, Y., Wei, Y., Hong-wei, M., Kang, T., … Zhan-sheng, J. (2017). Interferon-Induced Transmembrane Protein 3 Inhibits Hantaan Virus Infection, and Its Single Nucleotide Polymorphism rs12252 Influences the Severity of Hemorrhagic Fever with Renal Syndrome. Frontiers in Immunology, 7(535). doi:10.3389/fimmu.2016.00535

Xu, Q. F., Zheng, Y., Chen, J., Xu, X. Y., Gong, Z. J., Huang, Y. F., … Lai, W. (2016). Ultraviolet A Enhances Cathepsin L Expression and Activity via JNK Pathway in Human Dermal Fibroblasts. Chin Med J (Engl), 129(23), 2853–2860. doi:10.4103/0366-6999.194654

Zhang, Y. H., Zhao, Y., Li, N., Peng, Y. C., Giannoulatou, E., Jin, R. H., … Dong, T. (2013). Interferon-induced transmembrane protein-3 genetic variant rs12252-C is associated with severe influenza in Chinese individuals. Nat Commun, 4, 1418. doi:10.1038/ncomms2433

Zhao, X., Guo, F., Liu, F., Cuconati, A., Chang, J., Block, T. M., & Guo, J. T. (2014). Interferon induction of IFITM proteins promotes infection by human coronavirus OC43. Proc Natl Acad Sci U S A, 111(18), 6756–6761. doi:10.1073/pnas.1320856111

Zhao, X., Sehgal, M., Hou, Z., Cheng, J., Shu, S., Wu, S., … Guo, J. T. (2018). Identification of Residues Controlling Restriction versus Enhancing Activities of IFITM Proteins on Entry of Human Coronaviruses. J Virol, 92(6). doi:10.1128/JVI.01535-17

Zhao, X., Zheng, S., Chen, D., Zheng, M., Li, X., Li, G., … Guo, J. T. (2020). LY6E Restricts the Entry of Human Coronaviruses, Including the Currently Pandemic SARS-CoV-2. J Virol. doi:10.1128/JVI.00562-20

Zheng, M., Zhao, X., Zheng, S., Chen, D., Du, P., Li, X., … Lin, H. (2020). Bat SARS-Like WIV1 coronavirus uses the ACE2 of multiple animal species as receptor and evades IFITM3 restriction via TMPRSS2 activation of membrane fusion. Emerg Microbes Infect, 9(1), 1567–1579. doi:10.1080/22221751.2020.1787797

Zhou, P., Yang, X. L., Wang, X. G., Hu, B., Zhang, L., Zhang, W., … Shi, Z. L. (2020). A pneumonia outbreak associated with a new coronavirus of probable bat origin. Nature, 579(7798), 270–273. doi:10.1038/s41586-020-2012-7

Zhou, Z., Wang, R., Yang, X., Lu, X. Y., Zhang, Q., Wang, Y. L., … Wang, H. (2014). The cAMP-responsive element binding protein (CREB) transcription factor regulates furin expression during human trophoblast syncytialization. Placenta, 35(11), 907–918. doi:10.1016/j.placenta.2014.07.017

Zhu, N., Zhang, D., Wang, W., Li, X., Yang, B., Song, J., … Research, T. (2020). A Novel Coronavirus from Patients with Pneumonia in China, 2019. N Engl J Med, 382(8), 727–733. doi:10.1056/NEJMoa2001017

Zhu, Y., Feng, F., Hu, G., Wang, Y., Yu, Y., Zhu, Y., … Zhang, R. (2020). The S1/S2 boundary of SARS-CoV-2 spike protein modulates cell entry pathways and transmission. bioRxiv, 2020.2008.2025.266775. doi:10.1101/2020.08.25.266775

